# ImAge: an imaging approach to quantitate aging and rejuvenation

**DOI:** 10.1101/2022.10.16.512441

**Authors:** Martin Alvarez-Kuglen, Delany Rodriguez, Haodong Qin, Kenta Ninomiya, Lorenzo Fiengo, Chen Farhy, Wei-Mien Hsu, Aaron Havas, Gen-Sheng Feng, Amanda J. Roberts, Rozalyn M. Anderson, Manuel Serrano, Peter D. Adams, Tatyana O. Sharpee, Alexey V. Terskikh

**Affiliations:** Sanford Burnham Prebys, La Jolla CA 92037, USA; UCSD, Department of Physics, La Jolla, CA 92093, USA; Salk Institute for Biological Studies, La Jolla, CA 92037, USA; UCSD School of Medicine, 9500 Gilman Drive, La Jolla, CA 92093, USA; University of Wisconsin, Madison, WI 53705, USA; The Scripps Research Institute, La Jolla, CA 92037, USA; Institute for Research in Biomedicine (IRB Barcelona), Barcelona Institute of Science and Technology (BIST), Barcelona 08028, Spain; Altos Labs, Cambridge Institute of Science, Granta Park CB21 6GP, UK

## Abstract

Biomarkers of biological age that predict the risk of disease and expected lifespan better than chronological age are key to efficient and cost-effective healthcare^1–3^. To advance a personalized approach to healthcare, such biomarkers must perform on the individual rather than population level, demonstrate single cell resolution, and provide scalable and cost-effective measurements. We developed a novel approach – image-based chromatin and epigenetic age (ImAge), that utilizes image texture features based on the patterns of chromatin and epigenetic marks in single nuclei. We observed the emergence of intrinsic trajectories of ImAge using dimensionality reduction without regression on chronological age. ImAge was correlated with chronological age in all tissues and organs examined and was consistent with the expected acceleration and/or deceleration of biological age in chronologically identical mice treated with chemotherapy or following a caloric restriction regimen, respectively. ImAge from chronologically identical mice inversely correlated with their locomotor activity (greater activity for younger ImAge), consistent with the essential role of locomotion as an aging biomarker. Finally, we demonstrated that ImAge is reduced upon partial reprogramming in vivo following transient expression of OSKM cassette in the liver and skeletal muscles of old mice and validated the power of ImAge to assess the heterogeneity of reprogramming. We propose that ImAge represents the first-in-class individual-level biomarker of aging and rejuvenation with single-cell resolution.

## INTRODUCTION

With a steady increase in average lifespan and population aging^4, 5^, the accelerated adoption of biomarkers of aging (i.e. biological age) is needed. Such biomarkers should perform better, compared to chronological age, to quantitate disease risks and thus improve the effectiveness and quality of care in mid-life and old age while reducing associated cost^6–9^.

Biomarkers of aging include grip strength and gait^10^, frailty indices^11, 12^, frailty-based clocks^13^, metrics of the immune system^14, 15^, telomere length^16^, glycosylation readouts^17^, levels of cellular senescence^18^, and DNA methylation (DNAm) clocks. The fascinating discoveries of Hannum^19^ and Horvath^19, 20^ have identified CpG sites in human blood and other tissues with age-dependent changes in DNAm, enabling the development of DNAm clocks. Subsequently, multiple other combinations of CpGs / clocks have been established^21–25^. This approach has produced excellent chronological clocks tailored to specific cell types or pan tissues^26–29^. However, separating the biological components from the chronological components remains a chellenge^26, 30^. The vast majority of cellular biomarkers are obtained from the blood. These biomarkers correlate with mortality^31, 32^, predict lifespan^23^, and are sensitive to lifespan-altering interventions^33, 34^.

These biomarkers are population-level tools and effective estimators in large study cohorts not designed to reflect the individuality of the organism for which the deviation from the average is measured. This is important because comparing biological function of an individual, which is highly context-dependent, with a “golden mean” may be irrelevant for a particular individual^35–37^. Population-level biomarkers vary significantly at the individual level^38^ partly because their levels could be confounded by a pathology, for instance, an inflammatory response, rather than indicate functional or biological age^38^. In addition, most of these biomarkers measure averages over large numbers of cells (bulk measurements) and are not designed to quantitate widely assumed cell heterogeneity in organs/tissues, which can only be addressed at the single-cell level. Here, we develop the measurement of chromatin age in a single sell.

The loss of epigenetic information has been proposed to be a cause of mammalian aging^39^, and transient expression of Yamanaka’s OSKM factors^40^ is sufficient for shifting towards the younger state at least some readouts in organs and tissues of old mice^41, 42^. Cell identities defined by the global epigenetic state of individual cells ensure organismal homeostasis^43, 44^. Epigenetic changes may precede the cell identity change^45^; the new cell identity is associated with a new epigenetic state^46–48^. The alteration of epigenetic states is generally associated with cellular functional and phenotypic changes^45–49^, including aging^39, 50^. Epigenetic changes during aging result in altered epigenome and chromatin accessibility, aberrant gene expression, reactivation of transposable elements, and genomic instability^51, 52^.

Which specific epigenetic marks best convey age-dependent alterations is unclear, however, several studies linked aging to the loss of heterochromatin and alterations in global and local levels of H3K9me3, H3K27me3, H4K20me3, and H3K4me3^51, 53, 54^. The pattern of active enhancers, marked with a combination of H3K27ac & H3K4me1^55^, is also age-dependent^56, 57^. It is likely that information collectively encoded by the above epigenetic marks will be relevant to determining the progress of epigenetic and, perhaps, functional aging.

Several years ago, we pioneered microscopic imaging of epigenetic landscapes rooted in the analysis of chromatin topography in single cells^58^. We employed immunolabeling with antibodies specific for histone modifications (e.g. acetylation and methylation marks) and automated microscopy to capture cell-specific patterns using image texture analysis, resulting in multiparametric signatures of cellular states^58^. Here, we took advantage of this technique to develop an image-based chromatin and epigenetic age (ImAge), a fundamentally different approach to studying aging compared to DNAm clocks. We discovered the emergence of chromatin trajectories of aging as an intrinsic property of chromatin evolution with time. We observed a good correlation between ImAge and chronological age in mouse peripheral blood mononuclear cells (PBMCs) and several solid organs without regression on chronological age. Critically, ImAge of skeletal muscles from chronologically identical mice inversely correlated with their locomotor activity, suggesting its utility beyond correlation with chronological age. Finally, we documented the utility of ImAge to quantitate sample-wise and single cellwise heterogeneity of rejuvenation following partial reprogramming in vivo.

## RESULTS

### Microscopic imaging of chromatin and epigenetic determinants

We have pioneered the technique of microscopic imaging of chromatin and epigenetic determinants, which captures patterns of nuclear immunostaining of epigenetic marks, derives multiparametric signatures, and employs machine learning to identify multiparametric signatures of normal and neoplastic cell states^58^. We applied this technique first to investigate aging, focusing on freshly isolated primary cells (**Fig.1a**). We employed different combinations of histone modification marks with well-recognized roles in aging with multiplexing often limited by the availability of compatible primary antibodies. We use DAPI to label DNA/chromatin, which enables joint registration of all other epigenetic marks and provides chromatin pattern information. Acquired fluorescence images are processed using custom-built Python scripts, including nuclei segmentation with StarDist^59^, and feature extraction using threshold adjacency statistic (TAS)^60, 61^. It is important to note that, although the raw data are acquired from single cells (single nuclei), bootstrapped means (n=200 cells per bootstrap) were used to capture the properties of the single cell distribution while improving accuracy and equalizing the number of data points per sample. The size of the bootstraps was determined empirically by finding the lowest n while maximizing separation accuracy (**Extended Data Figs. 1** and **3**). We employed several dimensionality reduction techniques using Euclidean and non-linear, hyperbolic distance measures, multidimensional scaling (MDS), UMAP^62^, hSNE^63^, and hyperbolic MDS (HMDS)^64^, which are a modification of tSNE^65^ and MDS performed in the hyperbolic space.

**Fig. 1.**
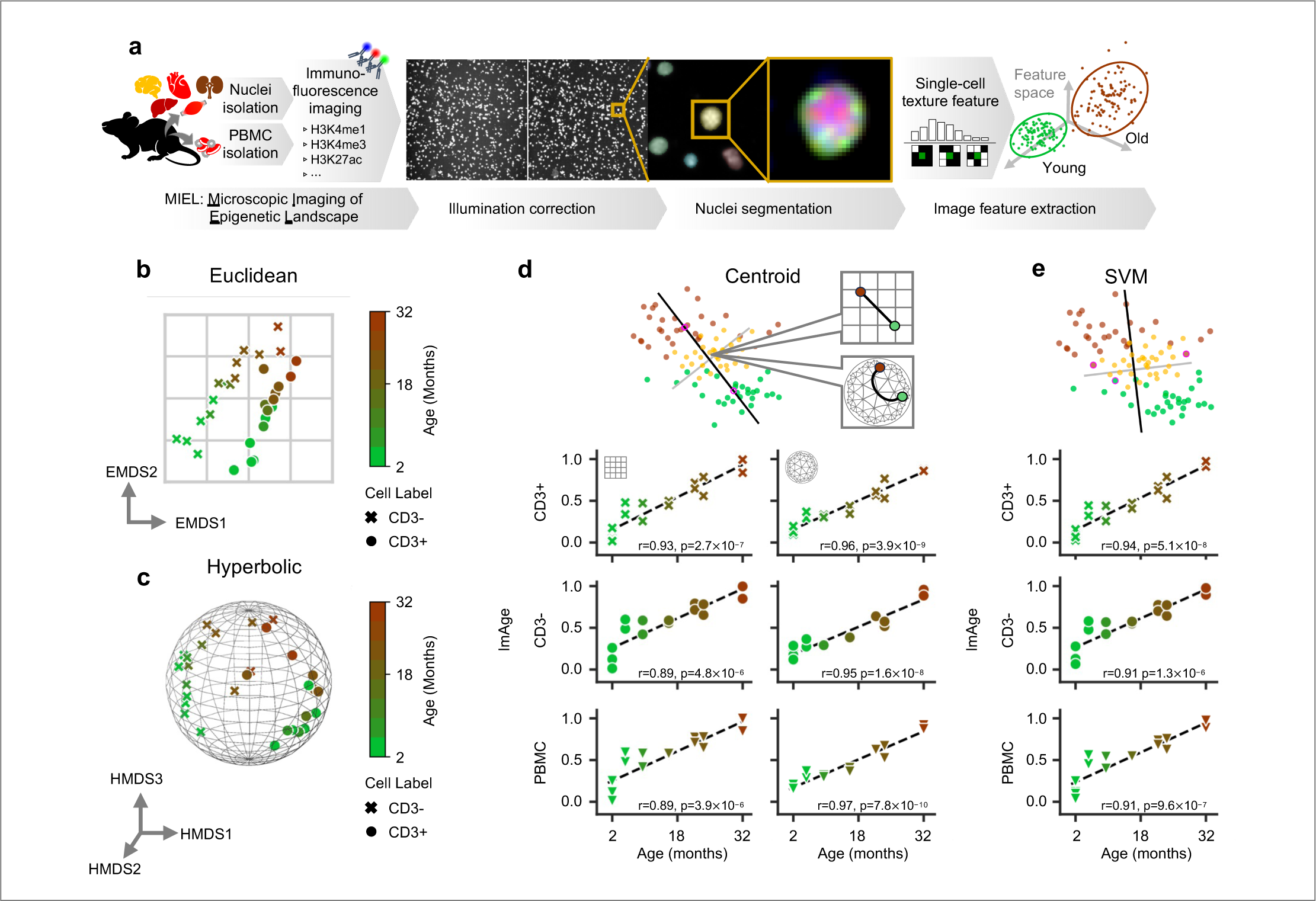
Emergence of chromatin trajectories of aging. **a**, Graphical representation of the MIEL workflow: Nuclear isolation, immunofluorescence imaging, image preprocessing, nuclear segmentation, texture feature extraction, and downstream analysis. **b-c**, ImAge calculations and regression analysis of CD3+ and CD3-subset of PBMCs from C57BL/6NJ males aged from 1.7 to 32 months (1.7, 2.2, 5.3, 8.7, 15.1, 21, 22.3, 32.2) (n=2 per age group). **b**, 2-dimensional EMDS of texture features. **c**, 3-dimensional HMDS of texture features. d-e, Graphical representation of the method of ImAge axis construction above scatterplots of the resulting ImAge measurements versus age for the CD3+ subset (top), the CD3-subset (middle), and the whole population (PBMC) (bottom). **d**, ImAge using the geodesic connecting the centroids of the youngest and oldest groups in Euclidean or hyperbolic space. **e**, ImAge using Linear SVM fit to the youngest and oldest groups. PBMC: peripheral blood mononuclear cells. E/HMDS: Euclidean/hyperbolic multi-dimensional scaling, respectively.

### The emergence of ImAge trajectories of aging

We assayed mouse PBMCs from C57BL/6NJ males aged from 1.7 to 32 months (1.7, 2.2, 5.3, 8.7, 15.1, 21, 22.3, 32.2 months). Cells were labeled with anti-H3K4me1, anti-CD3, and DAPI; signatures were computed using H3K4me1, anti-CD3 and DAPI channels as described before^58^. We employed metric Euclidean space (EMDS), to embed the data in a reduced 2-dimensional Euclidean space. EMDS revealed a clear separation of data points by cell type, CD3+ vs CD3-, mainly by the first dimension (EMDS1), and a clear age-related trajectory within each lineage, mainly by the second dimension (EMDS2) (**Fig. 1B**). Because EMDS aims to preserve distances when mapping from high to low dimensional space, we reasoned that the observed age-related progression should be a dominant characteristic of the geometry of the image feature space itself.

Given the high dimensional nature of this data, we sought out confirmation of the Euclidean results using a non-linear hyperbolic space, which has recently shown to be particularly helpful for analyzing high dimensional biological datasets^63, 66–68^. We sampled the CD3+/- single-cell data and embedded their relative distances into a 12 dimensional hyperbolic space with curvature (𝜅 = 7.2) via hyperbolic MDS (HMDS), optimized as previously described^63, 69^. We visualized this embedding using three-dimensional (3D) hyperbolic space and again observed a clear separation of epigenetic signatures between CD3+ and CD3-cells, as well as the emergence of age-related trajectory for both CD3+ and CD3-subsets of the PBMCs (**Fig. 1C**). These findings confirmed the emergence of intrinsic aging trajectories within epigenetic signatures of mouse CD3+/- subsets of PBMCs in two different geometries.

Thus, we aimed to find the simplest method to extract this trajectory, which we will henceforth call the ImAge axis and the progression along this axis as ImAge. To this end we first found the geodesic (shortest path) connecting the centroids of the youngest and the oldest mice in the full image feature space and found the projection of the data onto this geodesic, henceforth called the centroid based ImAge axis / ImAge (calculated separately for CD3+ cells, CD3-cells, and PBMCs). In the Euclidean space, we observed a strong correlation of centroid based ImAge with chronological age for CD3+ cells (Pearson-R = 0.92, p = 2.7×10^-7^), CD3-cells (Pearson-R = 0.88, 4.8×10^-6^), and total PBMCs (Pearson-R = 0.89, 3.9×10^-6^) (**Fig. 1d**, left). Likewise, we computed the geodesic in the hyperbolic space and projected the data onto that geodesic (calculated separately for CD3+ cells, CD3-cells, and PBMCs). Note that hyperbolic centroid-based ImAge provides stronger correlation with chronological age for CD3+ cells (Pearson-R = 0.96, p = 3.9×10^-9^), CD3-cells (Pearson-R = 0.95, 1.6×10^-8^), and total PBMCs (Pearson-R = 0.97, 7.8×10^-10^) (**Fig. 1d**, right).

Critically, both in Euclidean and hyperbolic spaces, the distance between the youngest and the oldest mice along the ImAge geodesic was greater than the orthogonal distances to the geodesic (**Extended Data Fig. 2**). This indicates that changes along the ImAge trajectory are a dominant source of changes in the dataset. Note that no regression on chronological age was necessary to reveal the ImAge trajectory: the trajectory is revealed as a dominant feature of the dataset geometry in both Euclidean and hyperbolic spaces, as seen in the EMDS and HMDS embeddings (**Fig. 1b** and **c**).

**Fig. 2.**
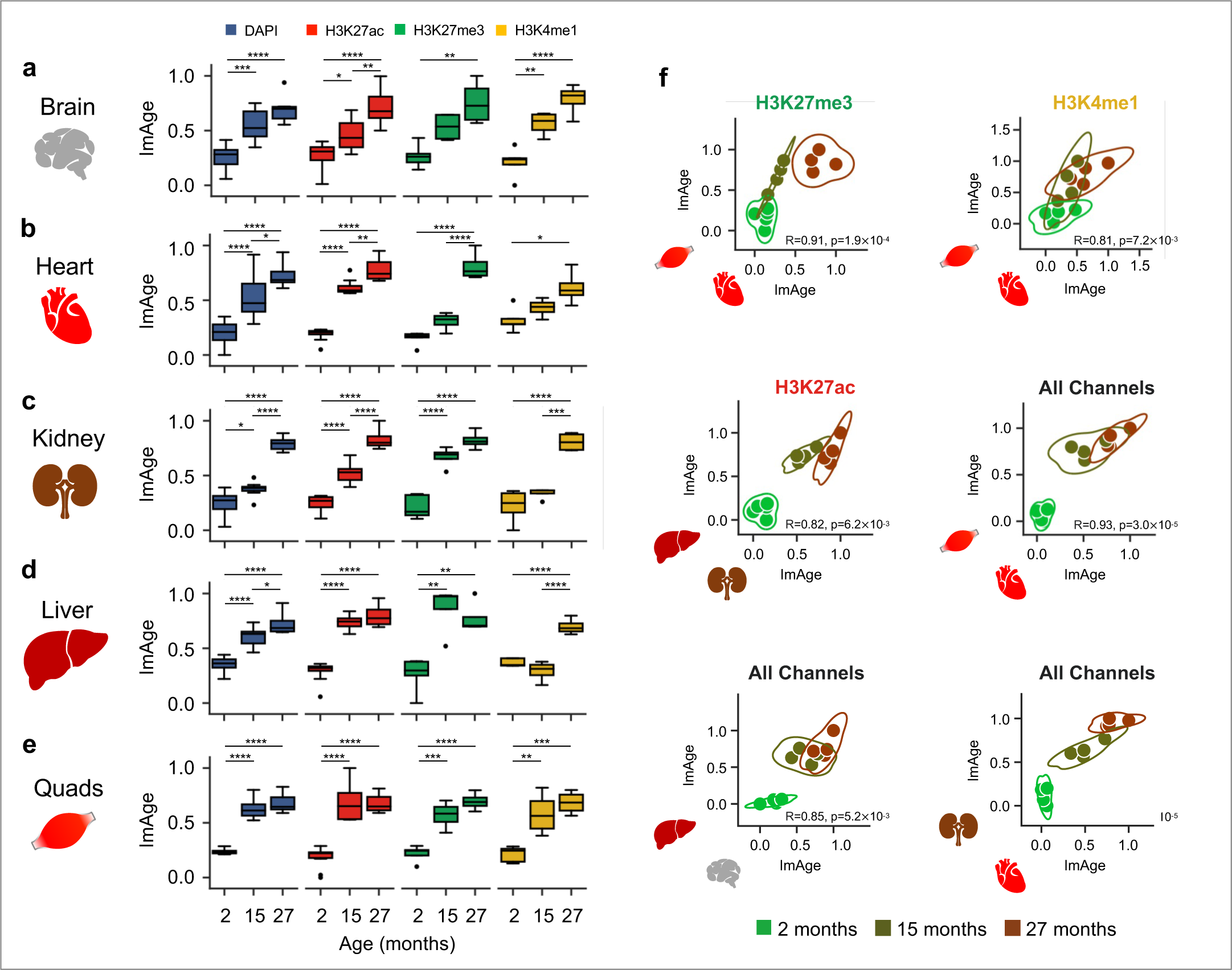
Age-related ImAge progression in multiple solid organs. **a-e** ImAge measurements and accuracy calculations on isolated nuclei from quadriceps (quads), liver, kidney, cardiac muscle (heart) and brain collected from three differentially aged cohorts of mice: 2 months (n=5) 15 months (n=4) and 27 months (n=4). Two plates were analyzed, both immunolabeled with H3K27ac+DAPI and then with either H3K27me3 or H3k4me1. Data for overlapping channels (DAPI & H3K27ac) were combined for computations. Boxplots min-max normalized 0-1 test set of bootstrapped data is shown. Differences of means were calculated via Tukey’s HSD. **f**, Consistent correlations were observed between skeletal muscle & heart (top row, bottom left), as well as brain & liver (bottom center), and heart & kidney (bottom right). Spearman’s R and p-values with Bonferroni correction for multiple comparisons. Significance values for all tests shown represent: * = p < 0.05, ** = p < 0.01, *** = p < 0.001, **** = p < 0.0001

To improve the robustness of the Euclidean method, we reasoned an approach to extract the geodesic which does not deform the feature space yet accounts for the potentially complex single-cell distribution was in order. Therefore, we utilized a linear support vector machine (SVM) to select the robust hyperplane separating the youngest and the oldest groups. The ImAge axis was then defined as the orthonormal vector to the hyperplane which passes through the region of highest density when the data (youngest and oldest groups) is projected to the hyperplane (**xrefFig. 1e**, top). As before, all intermediate time points were projected onto the ImAge axis without modifications (**Fig. 1d**). The correlation of chronological age and ImAge using SVM is intermediate to that obtained from the original Euclidean centroid-based ImAge and hyperbolic centroid based ImAge for CD3+ cells (Pearson-R = 0.94, p = 5.1×10^-6^), CD3-cells (Pearson-R = 0.91, 1.3×10^-8^), and total PBMCs (Pearson-R = 0.91, 9.6×10^-7^) (**Fig. 1e**) (**Extended Data Fig. 1**). The improvements seen with SVM is likely because of non-normality of the data distribution. Therefore, we will use SVM to construct ImAge axes on all subsequent organs and tissues when operating in a Euclidean space and will use the original centroid-based method to construct the ImAge axis when operating in hyperbolic space. Both will be henceforth referred to as ImAge and we will make clear which distance metric was used as needed.

### Comparative analysis of ImAge trajectories in solid organs

Recent studies suggested that organs and tissues may age at a different pace in the same organism^70, 71^. We investigated the distribution of ImAge readouts in 5 major organs, including brain, heart, kidney, liver, and skeletal muscles (quadriceps) in 3 cohorts of mice, young (2 months), middle age (15 months), and old (27 months). To directly compare ImAge trajectories in different organs and tissues, we developed a protocol to isolate nuclei from flash-frozen solid tissues (nuclei are PFA fixed immediately upon isolation to prevent any changes in chromatin state) and perform the analysis of chromatin and epigenetic landscape like that described for the freshly fixed cells (see Materials and Methods). We employed antibodies specific for H3K27me3, H3K27ac, H3K4me1 (+DAPI) to compute multiparametric signatures as previously described ^58^.

A Euclidean ImAge axis was then constructed between the youngest (2 months) and oldest (27 months) samples for each organ separately. We investigated the distribution of ImAge readouts in 5 major organs, including brain, heart, kidney, liver, and skeletal muscles (quadriceps) in 3 cohorts of mice, young (2 months), middle age (15 months), and old (27 months). In all tissues and organs analyzed, we observed that ImAge was increased with chronological age (**Fig. 2**). We observed monotonically increasing ImAge trajectories for all epigenetic marks and organs tested (**Fig. 2a-e**). Interestingly, the statistical significance appeared to be mark dependent. Separation accuracy was calculated per channel and for all channels combined (**Extended Data Fig. 3a-e**).

Given that 4 organs from the same mice were used per age group, we could compare the relative pace of ImAge progression in these organs. To select the most robust correlations we removed 1 age group iteratively and corrected for multiple comparisons (Bonferroni). We discovered a strong and robust correlation of ImAge in the heart and quads across several epigenetic marks and chronological age groups (Spearman-R: R=0.91, p=1.9×10^-4^, R=0.81, p=7.2×10^-3^, R=0.93, p=3.0×10^-5^ for H3K27me3, H3K4me1, and all channels combined, respectively) (**Fig. 2f**). Other pairs correlated when all marks were combined (liver & kidney, heart & brain), but were not consistent across individual marks (**Fig. 2f**). There are likely additional significant and robust correlations to be further substantiated with a larger dataset and sample size (**Extended Data Fig. 4**).

**Fig. 3.**
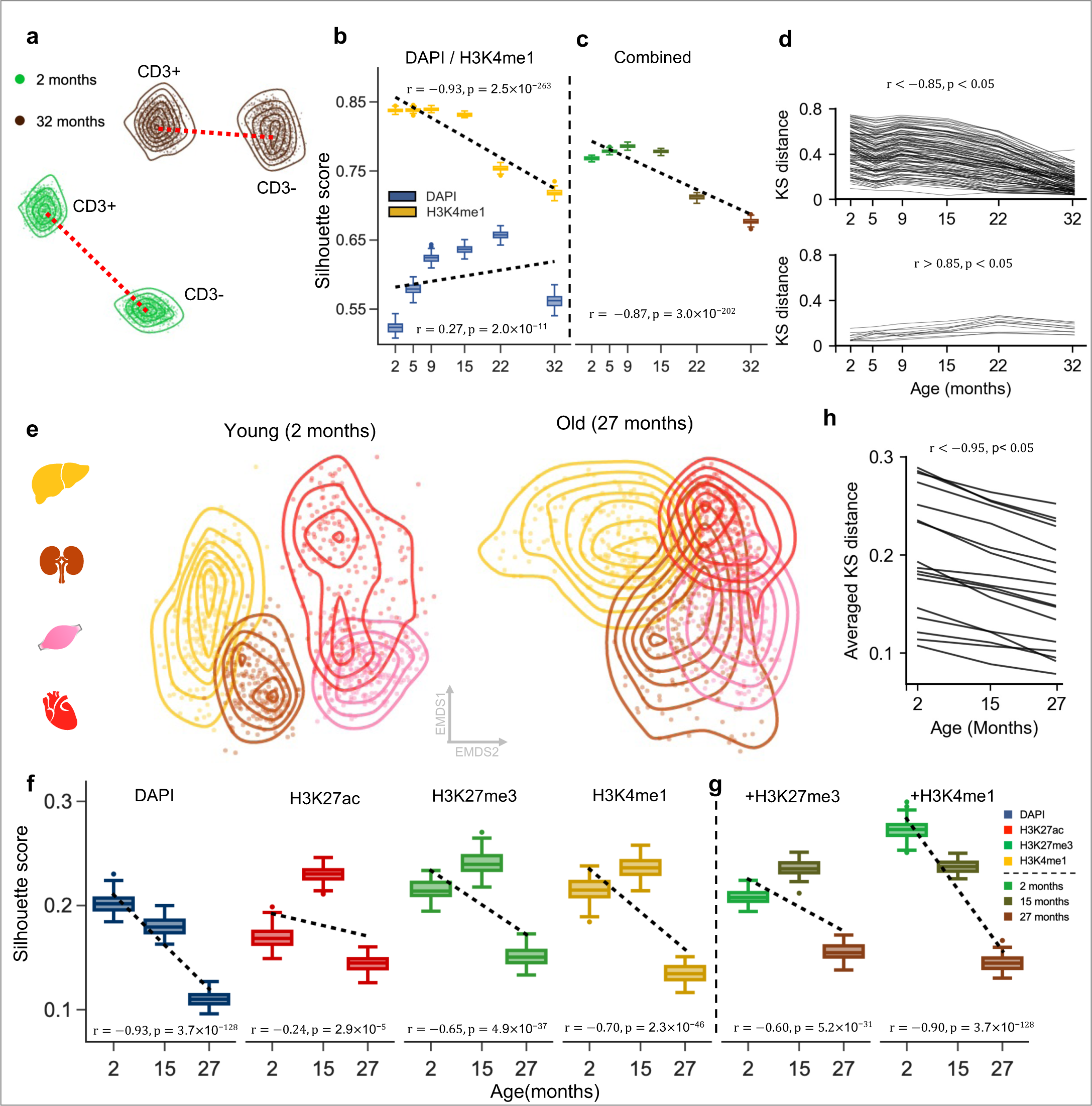
Age-related loss of cell type-specific chromatin and epigenetic information. **a**, A 2-dimensional EMDS of young (1.7 months) and old (32.2 months) CD3+ and CD3-subsets of PBMCs. **b, c**, Silhouette scores of CD3+ and CD3-subsets at indicated ages for individual marks (b) or their combination (c) using the information distance metric (based on mutual information and Shannon entropy, see Methods). **d**, the Kolmogo-rov-Smirnov (KS) distance analysis of CD3+ and CD3-subsets across indicated ages performed on significant features. **e**, A 2-dimensional EMDS of young (2 months) and old (27 months) liver, kidney, quads, and heart. **f**, **g**, Silhouette scores of 5 organs at indicated ages for individual marks (f) or their combination (g) using the information distance metric. **d**, the Kolmogorov-Smirnov (KS) distance analysis of 4 organs across indicated ages performed on significant features. Significant features were selected based on: 1) statistically significant (p < 0.05, Pearson |R|>0.85) KS distances between cell types and 2) a statistically significant (p < 0.05, Pearson |R|>0.95) correlation between KS distance and age.

**Fig. 4.**
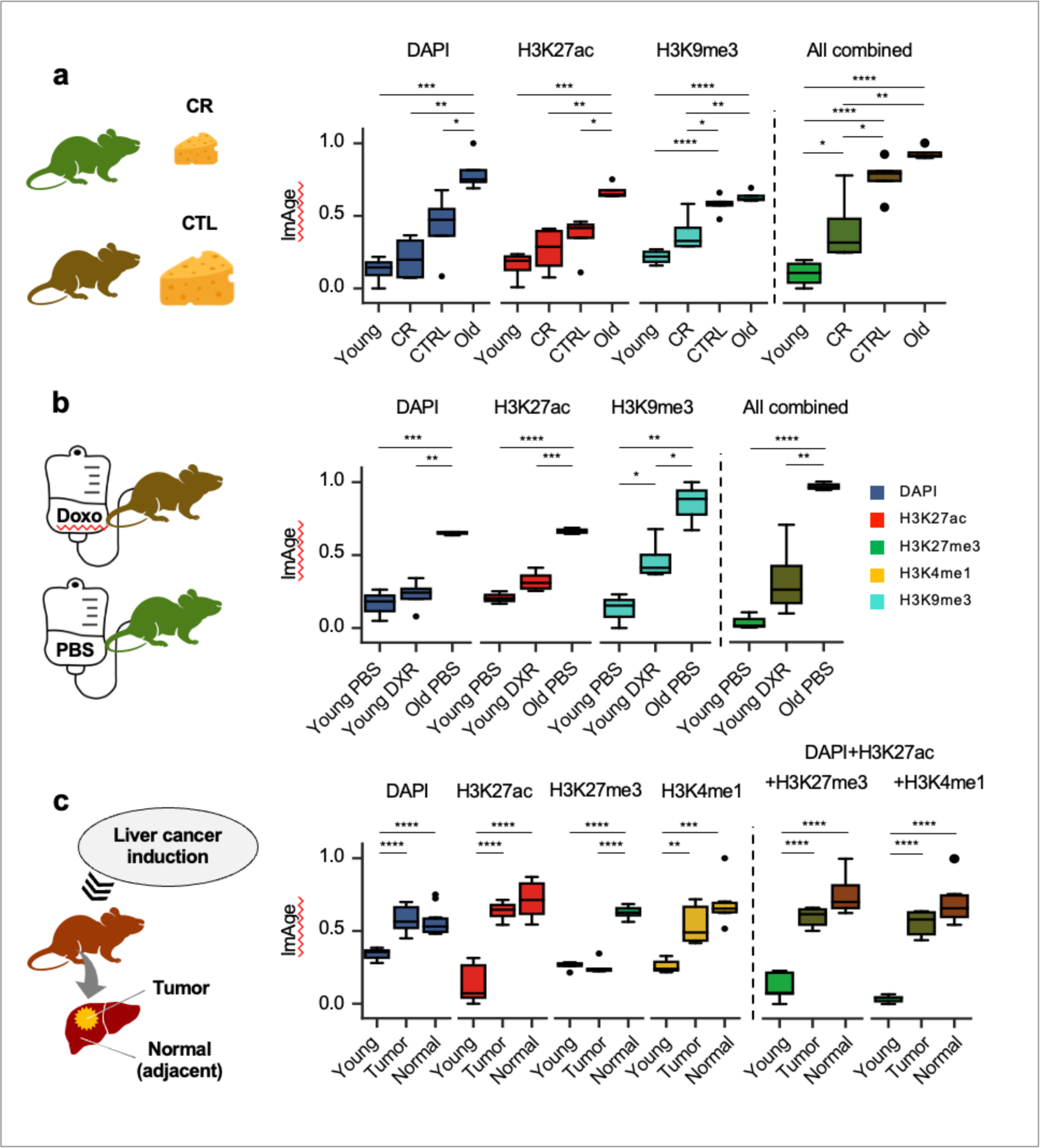
Diet, chemotherapy, and cancer affect ImAge. **a-c** ImAge calculations separated by epigenetic marks and several marks combined for indicated conditions for Caloric Restriction (CR), Doxorubicin (DXR), or induced hepatocarcinomas (tumor). **a**, young (3 mo., n=4) and old (24 mo., n=4) control mice were used to construct an ImAge axis upon which CR (7 mo., n=4) and control (7 mo., n=4) ImAge values were measured. **b**, young (1 mo., n=3) and old (27 mo., n=3) control mice treated with PBS were used to construct an ImAge axis upon which young DXR treated mice (1 mo., n=4) ImAge values were measured. **c**, liver tissue from old mice (8 mo., n=3) with induced tumors was separated by the presence or absence of tumors. Normal old tissue along with young control mice (2 mo., n=3) were used to construct an ImAge axis upon which old tumor ImAge values were measured. In (c), two plates were analyzed, both immunolabeled with H3K27ac+DAPI and then with either H3K27me3 or H3k4me1. Data for overlapping channels (DAPI & H3K27ac) were combined for computations. Significance was calculated using Tukey’s HSD. All ages shown are in months. p-value cutoffs are as follows: *: 0.01 < p <= 0.05; **: 0.001 < p <= 0.01; ****: p <= .0001

### Age-related erosion of cellular epigenetic and information identity

From our lower-dimensional visualization of ImAge trajectories using MDS in both Euclidian and hyperbolic spaces (**Fig. 1b** and **c**), it visually appears that the separation between T cells (CD3+) and non-T cells (CD3-) is reduced with age. To circumvent any bias attributed to different dimension reduction techniques, we quantified the age-related separation between CD3+ and CD3- and several tissues by computing both information distance using the silhouette score calculated across all features and the Kolmogorov-Smirnov distance (KS distance) calculated for each individual feature using the raw data (see materials and methods). The silhouette score computed across all features revealed an age-related decrease in information distance between the CD3+ (T cell) and CD3- cells within PBMC samples validating the visual assessment of MDS embeddings (**Fig. 3A, B**). Remarkably, we observed a similar age-related decrease in information distance across the liver, kidney, heart, and quads (**Fig. 3d** and **e**), while the brain remained separated from the other organs (**Extended Data Fig. 5**). Next, we calculated the KS distance across features significantly correlating with age (see materials and methods for details) and observed a prevailing negative correlation between the separation of cell/organ type and age, observed both for CD3+/CD3- cells (Pearson R < -0.85, p < 0.05) (**Fig. 3c**) and liver, kidney, heart, and quads (Pearson R < -0.95, p < 0.05) (**Fig. 3f**) for the vast majority of significant features. For the tissue samples, the number of significant features is comparatively fewer due to the presence of only three distinct chronological age groups available for calculating the correlation between KS distance and age. In sum, our analysis suggests an age-related erosion of cellular epigenetic identity and a loss of cell- and tissue type-specific information.

### ImAge distinguishes experimental perturbations affecting biological age

Previous work demonstrated the age-accelerating effect of widely used chemotherapeutic agents^72–74^. We followed Demaria et al., 2017 ^75^ protocol for doxorubicin treatment (10 mg/kg, i.p.; controls received PBS). Live hepatocytes were isolated 21 days post injection, to avoid acute stress response to DNA damage. Purified hepatocytes (2-step perfusion method^76^) were plated in 384 well plates, fixed, immunolabelled with anti-H3K9me3 and anti-H3K27ac (+DAPI), imaged, multiparametric signatures were computed as before ^58^ and projected on the ImAge axis as described above. We observed that doxorubicin treatment shifted the ImAge readouts from freshly isolated liver hepatocytes toward that of an older age (**Fig. 4a**). Statistical significance was driven by H3K9me3; trend for H3K27ac and DAPI. Multiple studies suggest that calorie restriction (CR) slows aging in diverse species ^77–79^. We compared the effect of CR on ImAge of liver hepatocytes. C57BL/6J males were fed ad libitum (control) or 75% of ad libitum (25% CR diet) from 2 months of age. Nuclei were purified from frozen liver tissues as previously described ^80^, distributed in 384 well plates, fixed, and immuno-labelled with anti-H3K9me3 and anti-H3K27ac (+DAPI). Images and multiparametric epigenetic signatures were acquired as before ^58^ and projected on the ImAge axis as described above. We observe that CR treatment, on average, shifts the ImAge readouts in liver hepatocytes towards that of a younger age (**Fig. 4b**). Statistical significance was driven by H3K9me3; with non-significant trends for H3K27ac and DAPI. Indeed, changes in H3K9me3 have been robustly associated with aging^52, 81–83^ further substantiating that H3K9me3 ImAge readouts in liver cells track with experimental perturbations of biological age.

### ImAge of liver tumors and adjacent normal tissue

We compared ImAge of liver tumors induced by diethylnitrosamine injection (postnatal day15) with normal tissues from the same livers in 8 months old C57BL/6NJ mice using immunolabeling for H3K27me3, H3K27ac, H3K4me1, and DAPI. No difference in ImAge was observed using DAPI or H3K27ac, whereas tumors appeared significantly younger with H3K27me3 (p <= 0.0001, Tukey’s HSD), and a trend was observed for H3K4me1(**Fig. 4c**) (**Extended Data Fig. 5**).

**Fig. 5.**
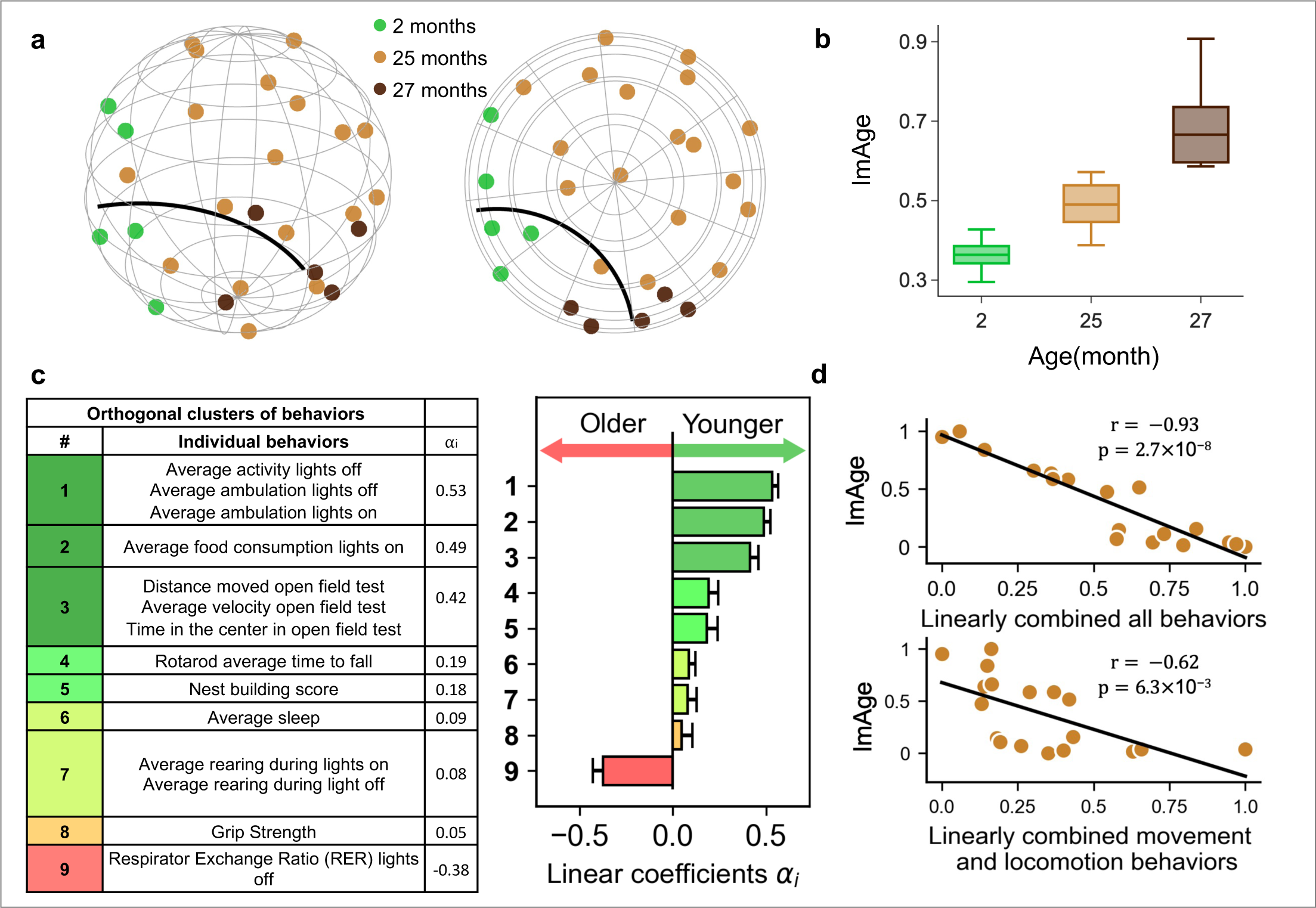
Locomotor activity is a salient correlate of ImAge in chronologically identical mice. **a**, a 3-dimensional representation of Hyperbolic embedding (HDMS) and its 2-dimensional projection (view from the top) of the young (2 months) and old (27 months) mouse quadriceps samples utilized as references to obtained centroids for the ImAge axis. Quadriceps samples from chronologically identical (25 months) mice from the behavioral cohort were co-embedded with reference mice to obtain (**b**) their ImAge distribution between the reference samples. **c**, the 9 orthogonal clusters of behavior with the coefficients for linear optimization correlating behavioral/functional readouts and ImAge. The direction of ImAge association with each cluster (older, younger) is proportional to the cluster’s ⍺i. **d**, Correlations between ImAge and a linear combination of all behavioral readouts (top) and locomotor activities only, clusters 1, 3, 4, and 7 (bottom).

### ImAge correlates with age-related behaviors in chronologically identical mice

Next we investigated whether or not ImAge readouts map to functionally meaningful organismal readouts (which is to say, biological age) in chronologically identical samples. To this end, we conducted metabolic, cognitive, and motor behavioral tests in chronologically identical C57BL/6 males (25 months old, n=18) and acquired ImAge signatures from the skeletal muscles. We used young (2 months old, n=5) and old (27 months old, n=5) mice to derive the ImAge axis of chronological young and old controls. We focused on skeletal muscles (quadriceps), which bear the functional load of motor behavior and mediate systemic metabolism^84^. Nuclei were isolated from flash-frozen tissues, immunolabeled with H3K27ac + H3K27me3 antibodies (+DAPI), imaged, and multiparametric epigenetic signatures were acquired as previously described^58^.

We embedded the data points, which are bootstrap means of 200 samples, into a significantly reduced 9-dimensional hyperbolic space using information distance as the metric. The curvature of this hyperbolic space was determined to be -8.4 (see materials and methods). The dimensionality and curvature were optimized as previously described^63, 69^. Within this hyperbolic space, the shortest path connecting two data points is commonly referred to as a geodesic, which is curved instead of a straight line in the Euclidean space. We computed the geodesic between the centroids of young and old reference mice in the hyperbolic space and projected the coordinates of all mice onto this geodesic, resulting in a distribution of ImAge readouts. As with the other tissues, we observed a robust separation between the ImAge from young and old reference mice, and the ImAge readouts of experimental mice were distributed in between (**Fig. 5a** and **b**). To extract relationships between ImAge readouts and behavioral readouts, we used linear regression. To ensure we extracted only the most stable relationships, we excluded non-significant, highly variable behavioral readouts (see materials and methods). To reduce collinearity in the regression analysis we clustered correlated behaviors (**Extended Data Fig. 7 a-b**), thus deriving regression coefficients for 9 orthogonal clusters (**Fig. 5c**, see materials and method for details). The linear coefficient 𝛼_!_ is proportional to the contribution, positive or negative, of each cluster to ImAge. Because individual readouts are normalized and colinear within each cluster, we consider each of individual readouts within the cluster contribute to ImAge with the same coefficient 𝛼*_i_* Such computations enabled us to identify key metabolic readouts and behaviors with positive or negative contributions to ImAge (**Fig. 5c**).

The linear combination of the 9 orthogonal clusters of metabolic and behavioral readouts provided excellent (Pearson r=-0.93 p=2.7×10^-8^) correlation with ImAge (**Fig. 5d**, top). This strong and highly significant correlation of ImAge with a relatively small number of whole organism metabolic and behavioral readouts underscores the potential utility of ImAge as a single biomarker of functional or biological age in skeletal muscles and, possibly, other tissues and organs. The linear coefficient 𝛼_!_ is proportional to the contribution, positive or negative, of each cluster to ImAge. Because individual readouts are normalized and colinear within each cluster, we consider each of individual readouts within the cluster contribute to ImAge with the same coefficient 𝛼*_i_*. Such computations enabled us to identify key metabolic readouts and behaviors with positive or negative contributions to ImAge (**Fig. 5C**). Individually, cluster 2 and 3 significantly correlate with ImAge (**Extended Data Fig. 7 C**). Consistent with numerous previous studies, several forms of locomotor activity (cluster 1, 3, 4, 7) were negatively correlated with ImAge (greater activity for younger ImAge). Taken together these 4 major clusters provided strong (Pearson r=-0.62 p=6.3 E-3) correlation with ImAge (**Fig. 5D**, bottom). Respiratory Exchange Ratio (RER) during the dark phase (lights off) was positively correlated with ImAge (greater RER for older ImAge), in agreement with a recent cross-sectional study.

This linear combination of behavioral parameters can be conceptually framed as a weighted metric of behavioral performance. Notably, it reveals a substantial negative correlation with ImAge, signifying that as the mouse undergoes aging, there is a discernible decline in behavioral performance. Our findings suggest that ImAge integrates several behavioral/functional readouts, underscoring its utility beyond correlation with chronological age. We identified locomotor activity as the most salient behavior parameter that is higher for younger chromatin states. Critically, the ImAge readouts are derived from individual measurements and can be directly compared to each other (not the average), thus representing individualized biomarkers of chromatin age.

### ImAge reveals heterogeneity of in vivo reprogramming with OSKM factors

We inquired whether ImAge reports the reversion of the aging process using OSKM-driven partial reprogramming in vivo^41, 42^. We analyzed liver, heart, and skeletal muscle tissues (kindly provided by Dr. Serrano) from 13.8 month old i4F and intermate control mice treated for one week with a low dose of doxycycline (0.2 mg/ml)^42^. We focused on the available liver, which presented some reversion of the epigenomic changes associated with aging^42^, as well as the skeletal muscle.

Nuclei were isolated from the frozen liver and muscle samples of young (3.2 months), old (13.8 months), and old treated with doxycycline to overexpress OSKM factors (old-OSKM) (**Fig. 6a**, see ref. ^42^ for details of mouse samples and Materials and Methods for sample preparation). Nuclei were immunolabelled with H3K9ac, H3K27ac, H3K27me3, and DAPI, and confocal images were acquired with Opera Phenix (PerkinElmer). We obtained ImAge based on the distance from the hyperplane of a linear SVM using 3D TAS features extracted from confocal images. To compare each mouse at the individual level, the SVM model was trained using averaged data points obtained by bootstrapping 200 single-cell data points (see Methods for details of bootstrap sampling). The accuracy of the separation of young and old samples was 1.000 and 0.997 in testing for the liver and muscle samples, respectively, in 100 iterations of tests using random data splits (see Methods for details of training/testing scheme). There were significant differences (Mann-Whitney U-test, p < 0.05) between the median ImAges of young and old samples in both tissues (**Fig. 6b** and **c**). We observed that the mean ImAge in the old-OSKM group was significantly decreased compared to that in the old group, but was significantly higher than that of the young samples. These results suggest liver and muscle cells in old-OSKM mice are partially reprogrammed on average.

**Fig. 6.**
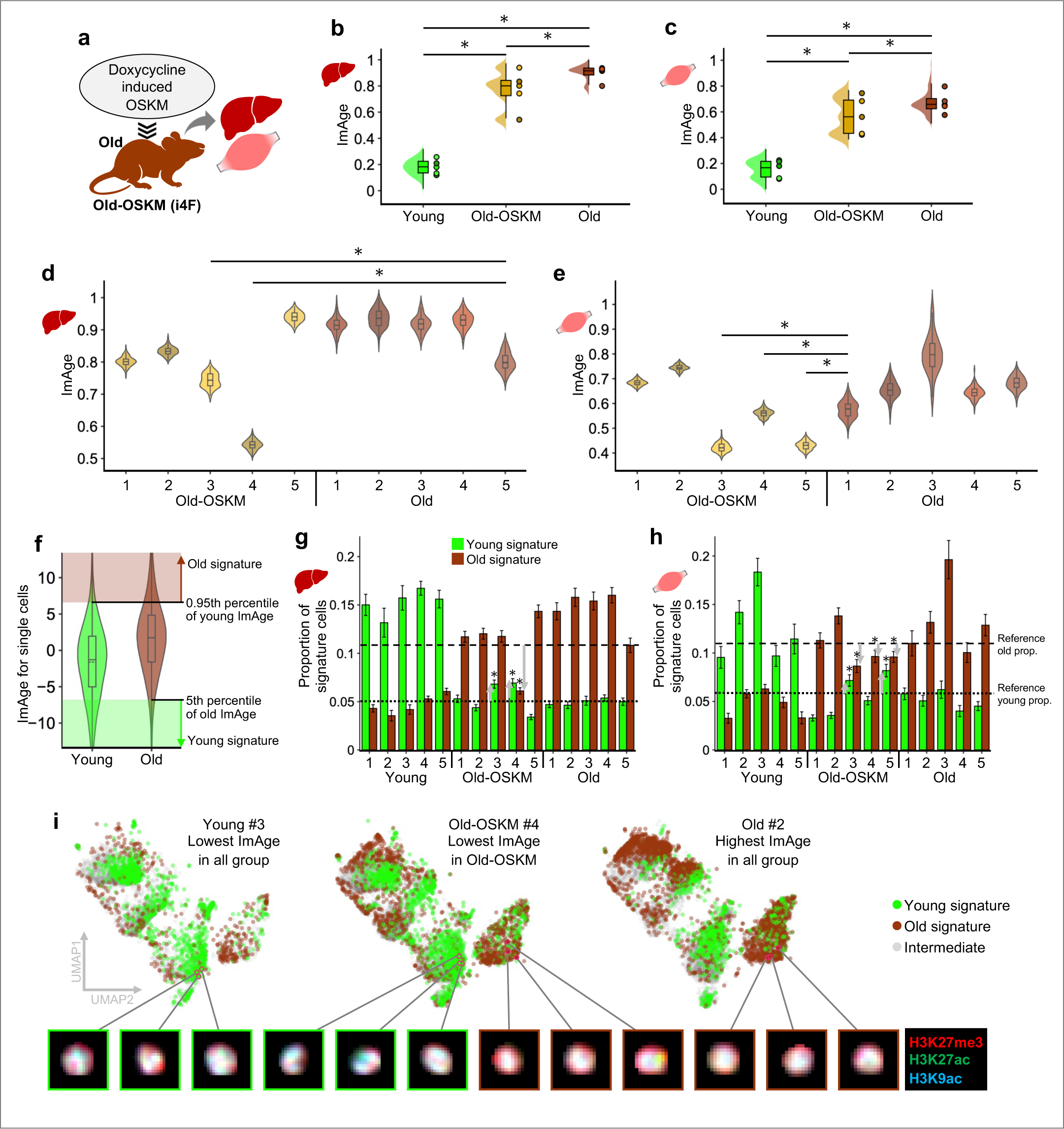
ImAge revealed heterogeneity of partial reprogramming in vivo. The chronological ages of young and old mice are 3.2 and 13.8 months, respectively. **a**, Evaluating the degree of reprogramming after doxycy-cline-induced OSKM factors (i4F mice, old-OSKM) in the liver and muscle. **b, c**, Distribution of ImAge (100 iterations of the test is shown) in the liver (b) and muscle (c). Dots on the right side of violin plots are the mean ImAge of individual mouse samples. **d, e**, Violin plots representing the distribution of the ImAge within individual samples in the liver (d) and muscle (e). Statistically significant differences were assessed between all old-OSKM samples and the old mice sample with the lowest ImAge (the “youngest” old mouse). **f**, Distribution of ImAge at a single-cell level. The mean accuracy of the young and old segregation was 0.620±0.001 (100 iterations). The young and old signatures are defined by the threshold of ImAge readout at 5th (and lower) and 95th (and higher) percentile values of old and young single cell ImAge readout, respectively. **g, h**, The proportion of young/old ImAge signatures in the liver (g) and muscle (h) for each animal. The gray arrows indicate an increase in young and a decrease in old signatures compared to the “youngest” old mouse defined in comparison of ImAge distributions (d and e). The dotted and dashed lines indicate the reference proportions of young and old signatures, respectively, in the “youngest” old mouse. i, Uniform manifold approximation and projection (UMAP) of single-cell texture features obtained from the liver samples. The green and brown data points represent the single cells with young and old ImAge signatures, respectively. The intermediate single cells are in grey. p<0.05 Mann-Whitney U-test.

Mouse sample-wise heterogeneity in response to the reprogramming was observed in the comparison of age group-wise variance and sample-wise ImAge readout distributions. At age group comparison, the variance of ImAge in the old-OSKM group was significantly higher than that of young and old groups (**Fig. 6b** and **c**, Levene’s test, p < 0.05). In the liver sample, there were significant differences (Mann-Whitney U-test, p < 0.05) between the median ImAges of two (#3 and #4) among the five old-OSKM mice samples as compared to the mouse with the lowest median ImAge in the old group (old #5) (**Fig. 6d**). In the skeletal muscle, we observe significant changes of ImAge in three old-OSKM mice (#3, 4 and 5) (**Fig. 6e**). Almost no changes were detected in the heart consistent with the phenotypic observations^42^ (**Extended Data Fig. 8d**). These findings suggest that ImAge detects cellular rejuvenation and reports variable reprogramming efficiency, at least in the liver and muscle, between individual organisms.

We observed different degrees of partial reprogramming per epigenetic marks/channels. To evaluate the channel-wise difference, the ImAge axis was constructed in the same way as described above for each channel (H3K9ac, H3K27ac, H3K27me3, and DAPI). In the liver, strong partial reprogramming was observed using DAPI and H3K9ac (**Extended Data Fig. 9a**), where OSKM-old #1-4 showed a significant decrease in ImAge. In H3K27me3, OSKM-old mice #1, #3, and #4 were found to be significantly reprogrammed. H3K27ac showed the weakest impact; only one significantly reprogrammed mouse was found (OSKM-old #4). In the muscle, all channels showed almost equal strength of reprogramming with significant decreases of ImAge observed in two mice (OSKM-old #5 and #3/#4), except for H3K9ac, which showed a significant decrease in three mice (OSKM-old #3-5). DAPI and H3K27me3 had the same mice pair significantly reprogrammed (#3 and 5).

We leveraged single-cell-nature of imaging, to get insights into the heterogeneity of reprogramming, which is of critical importance for rejuvenation^85^. We defined signature single-cells of the young and old groups and assessed the changes in the proportion of signatures in the old-OSKM group. We obtained ImAge readouts for single cells by projecting single-cell features on ImAge axis constructed on binned data points from 200 cells. Because of the heterogeneity of single cells, the tails of ImAge distributions were stretched, and the ImAge readouts for young and old groups were largely overlapped (**Fig. 6f**). In other words, at the single cell level, the boundary between young and old was not clear (accuracy: 0.62±0.001 in 100 iterations of tests).

Hence, the signature cells for each age group (young or old) were identified as single cells where the ImAge readout was hardly observed in the other group. For instance, young signature cells were cells with an ImAge readout lower than the 5th percentile of old ImAge, meaning young cells can only be observed at a maximum of 5 percent in the old group. Conversely, the old signature cells were determined by ImAge readouts higher than the 95th percentile of the young group (**Fig. 6f**, see Materials and Methods for details in determining the signature cells). We observed a significant increase in young signature cells and a decrease in old signature cells in the old-OSKM group on average (**Extended Data Fig. 9b**) in the liver and a significant decrease of the old signatures in the muscle (**Extended Data Fig. 9c**). We observed three different patterns of significant changes in the proportion of signature cells; an increase of young signature cells, a decrease of old signature cells and both of them. Particularly in the old-OSKM#4 in the liver, #3 and #5, which were three of the most reprogrammed mice, showed a significant increment of young signature and decrement of old signature (**Fig. 6g** and **h**). Because ImAge is rooted in the single-cell image analysis, cell nuclei with corresponding ImAge signatures can be identified (**Fig. 6i** and **Extended Data Fig. 10**).

## DISCUSSION

One of our most surprising findings is the discovery of intrinsic aging trajectories formed by multiparametric epigenetic signatures that could be rendered and visualized using Euclidian and hyperbolic metrics. Remarkably, no regression was required for ImAge axis construction, yet all intermediate age groups were naturally arranged in a highly ordered manner. The ImAge axis emerged (not enforced) as a dominant source of information in the image feature space (feature manifold), especially when this information is embedded in hyperbolic space. This suggests that we are extracting information related to the robust spatial organization of chromatin topography in individual nuclei and their evolution over time, which is biologically meaningful.

As an imaging-based approach, ImAge is complementary to previous aging biomarkers, such as DNAm clocks, which utilize a fundamentally different technique (sequencing). While ImAge is intimately linked with structural information and is blind to sequence level modifications, DNAm presents a sequence level approach which is largely blind to structural information. Another notable difference between ImAge and DNAm clocks is the utilization of linear regression: DNAm clocks universally regress CpG methylation levels against chronological age (i.e. building DNAm clock) to construct robust population level tools useful for numerous applications^22, 86^. DNAm clocks always select only a small subset of all CpGs in the genome (up to 1000 out of ∼2.8M, less than 0.1%); in contrast, ImAge captures the major characteristic of the data manifold. The highly ordered nature of the image feature space results in our ability to perform direct and precise measurements of the relative distances between individual samples.

However, these advantages do not come without limitation. DNAm clocks have become the gold standard for aging biomarkers, in part because they enable individual measurements to be easily compared against the population without reconstruction of entire model. A key part of the universality of these clocks is the stability of both DNAm itself and the biochemical technique to sequence CpGs. However, imaging is quite the opposite: image features are very sensitive to imaging conditions (such as microscope setup and antibodies), leading to many opportunities for instability of feature values between experiments. Currently, this limitation demands that ImAge axes must be computed for each new sample combination. However, it is worth noting that the spatial organization of chromatin, the imaging substrate, although fundamental variable^87^ is likely not the issue. Rather the pixel values per channel and their relative intensities (from differences in antibody concentration, laser intensity, detector, etc…). We hypothesize that the stability of the image-feature manifold itself is consistent across various imaging conditions. Future work should investigate the stability of this manifold across imaging conditions to separate biological variability at the single cell level from technical limitations. Fiduciary samples applied across multiple experiments and computational developments in deep learning provide empirical path to addressing these questions.

Single-cell DNA methylation analysis is challenging^88^ and may not be practical in the near future. While the image analysis presented here (20X magnification, mainly 2-dimensional maximum projection, TAS features) struggles to fully resolve age-related chromatin changes at the single-cell level, we have only scraped the surface of pre-existing advancements in microscopy and imaging analysis: high resolution confocal microscopy and advanced colocalization features present clear path towards extracting significantly more information from nuclear images and, potentially, highly structured single-cell resolution. Indeed our accuracy measurements show near perfect separation accuracy at a single-cell resolution is not far away (**Extended data Figs. 2**,**3**,**5**). Thus ImAge, being in its nescient stages of development, promises significant room for improvement on both imaging and computational fronts.

Imputing organ aging from plasma protein suggested different rates of organs and systems aging within the same organism (aging ageotypes)^71^ while the Tabula Muris Senis studies demonstrated a similar yet asynchronous inter-organ progression of aging^89^. Some degree of organ connectivity is plausible so that the decline of one organ can promote the dysfunction of other organs, accelerating organismal aging^90^. We observed an overall monotonic progression of ImAge along chronological age for all epigenetic marks and 5 organs tested. Together with our observations in PBMCs, it suggests that a monotonic correlation of ImAge and chronological age is universal. We observed a strong correlation of ImAge between the heart and skeletal muscles (quads) for all epigenetic marks (and DAPI) as well as the heart and kidney for all mark combinations. ImAge of the liver also correlated with the brain and kidney. Note that ImAge correlation between organs doesn’t mean that the pace of ImAge progression is the same in those organs. Incidentally, a metabolic imbalance of lipids accelerates inflammation and leads to lipotoxicity, mostly afflicting the kidney, heart, and skeletal muscle^91^, and lipotoxic insults precipitate aging^92^. Immobilization and atrazine induce oxidative stress in the liver, kidney, and brain, with curcumin and quercetin exhibiting a protective effect^93, 94^.

Cellular epigenetic identity is integral to governing cell functionality and maintaining tissue homeostasis. We observed an age-related juxtaposition of cellular states and a diminished separation of cellular identity among diverse tissues or cells, suggesting an erosion of cellular chromatin and epigenetic identity. Although we documented this effect for CD3+ and CD3- cells in the blood and between 5 different organs, this is likely a general phenomenon. This could be explained by the age-related erosion of epigenetic and chromatin landscape^39^ which increases the noise in part through cell-to-cell variation in gene expression^95^, loss of lineage fidelity with age, and activation of lineage-inappropriate genes during aging^81, 96–99^. Reactivation of human endogenous retroviruses during aging could also contribute to this process^100, 101^. Our observations are consistent with the information theory of aging, which proposes that aging in eukaryotes is associated with loss of epigenetic information over time^102–104^ and that loss could be causative^39^.

DNA damage is known to induce cancer and accelerate aging ^105–107^ and chemotherapy treatments have damaging effects on the entire organism and accelerate the aging process ^75, 108–110^ including increased frailty, chronic organ dysfunction, increase in cardiovascular diseases, cognitive impairment, and secondary cancers ^73, 111–114^. We observed a shift of ImAge towards old age in mice treated with DOX, resembling the observations in humans ^74^. The changes were driven by H3K9me3, suggesting possible changes in heterochromatin. If ImAge could faithfully report the effects of other chemotherapy drugs and in different organs, it will be useful to design interventions that increase the resilience to chemotherapy ^115^, and to reduce the aging effects of chemotherapy.

Because CR robustly increases maximum lifespan and delays biological aging in diverse scenarios ^77–79^, albeit the full picture could be more complex ^116, 117^. successfully applied CR regimens help to understand the biology of aging ^118–120^ and could help recovery after chemotherapy in clinic ^115^. We have observed a shift in ChromaScan readouts in CR animals towards that of young animals consistent with the phenotypic observations. It will be important to further refine these analyses, covering different CR regimens in mice and other species (e.g. monkeys), and to correlate the epigenetic changes with functional readouts at the organismal level as well as with various OMICs datasets (e.g. gene expression, chromatin accessibility, and enrichment for various epigenetic marks).

We observed younger ImAge in liver tumors, but only for H3K27me3 mark. Early studies suggested DNAm age acceleration in tumors compared to normal tissues^19, 20, 29^. However, recent studies matching tumor and normal tissue from the same organ/individual painted a more complex picture with deceleration of DNAm age in stomach adenocarcinoma^121^ and no change in triple-negative breast tumors^122^. Analysis of ∼439 endometrial cancers suggested unchanged DNAm age in 90% of tumors while DNAm age deceleration was associated with advanced diseases and shorter patient survival^123^. Given that extended partial reprogramming in vivo results in tumors^124^, a younger chromatin and epigenetic age of tumors is rather intuitive.

The largest group of behaviors that inversely correlated with ImAge comprised various flavors of locomotor activity. Age-related decline in locomotor activity is a universal feature of living organisms well documented in Drosophila^125^ and insects in general^126^, rodents^127^, dogs^128^, primates^129^, and humans^130^ where it is associated with cognitive impairment and decline in brain dopamine activity^131^. Locomotion is a very reliable factor for exploratory behavior in mice^132, 133^. The only correlate of older ImAge was the respiratory exchange ratio (RER), previously observed to be increased in old mice only in the dark phase^134^.

Remarkably, ImAge faithfully reported liver and muscle rejuvenation following one low dose cycle (one week of low dose in drinking water) of OSKM-mediated partial reprogramming in vivo, including the heterogeneity of rejuvenation at the individual organism and at the single cell level. The advantage of analyzing a single cycle is that we detect the “proximal” effects of OSKM overexpression. With multiple cycles, the combined effect over months may include many secondary effects. Although the final outcome is more pronounced, the underlying mechanism is likely to be “buried” in many layers of effects over months. Curiously, different mice appeared to be mostly reprogrammed for liver and skeletal muscles, with DAPI and H3K9ac being the most informative to detect the change. Although we didn’t have H3K9me3 for this experiment the combination of DAPI and H3K9ac (complimentary to H3K9me3) suggests that we might be detecting heterochromatin changes. It will be informative to directly compare the performance of DNAm clocks and ImAge to quantify the apparent heterogeneity of in vivo reprogramming^135^.

Given the robust correlation of ImAge with key physiological, behavioral, and metabolic metrics underlying functional decline with age, we posit that ImAge could be used as an individualized biomarker of functional age. It will be informative to analyze longitudinal ImAge trajectories in individual organisms. Individual ImAge trajectories are likely defined by the organism’s genetic makeup and the cumulative chromatin and epigenetic changes in response to environmental factors.

## MATERIALS AND METHODS

### Mice

Experiments were conducted according to guidelines and protocols approved by the Institutional Animal Care and Use Committee (IACUC) of Sanford Burnham Prebys Medical Discovery Institute. Data presented within this manuscript were obtained using male mice. C57BL/6 mice, ranging in age from 2 to 27 months old, and were obtained from National Institute on Aging, Aged Rodent Colonies (RRID:SCR_007317).

### Mouse behavioral studies

#### EchoMRI testing

The EchoMRI 3-in-1 instrument (EchoMRI LLC, Houston, TX) is a quantitative nuclear magnetic resonance (qNMR) imaging system for whole body composition analysis of unaesthesized small animals ^136, 137^, and qNMR body composition analysis with EchoMRI instrumentation has been proposed to be “gold standard” methodology for metabolic studies in the mouse ^138^. Following calibration, each mouse was put in a holder and placed into the EchoMRI chamber and lean mass, fat mass and water mass was calculated.

#### Optomotor test

The optomotor allows for assessment of visual ability and consists of a stationary elevated platform surrounded by a drum with black and white striped walls. Each mouse is placed on the platform to habituate for 1 minute and then the drum rotates at 2rpm in one direction for 1 minute, is stopped for 30 sec, and then rotates in the other direction for 1 minute. The number of head tracks (15 degree movements at speed of drum) is recorded. Blind mice do not track the moving stripes.

#### Comprehensive Laboratory Animal Monitoring System (CLAMS)

Indirect calorimetry was performed in acclimated, singly-housed mice using a computer-controlled, open-circuit system (Oxymax System) that is part of an integrated Comprehensive Lab Animal Monitoring System (CLAMS; Columbus Instruments, Columbus, OH: ^139, 140^). Testing occurred in clear respiratory chambers (20 × 10 × 12.5 cm) equipped with a sipper tube delivering water, food tray connected to a balance, and 16 photobeams situated in rows at 0.5 in intervals to detect motor activity along the *x*- and *z*-axes. Room air is passed through chambers at a flow rate of 0.5 L/min. Exhaust air from each chamber is sampled at 15-min intervals for 1 min. Sample air is sequentially passed through O2 and CO2 sensors (Columbus Instruments) for determination of O2 and CO2 content, from which measures of oxygen consumption (VO2) and carbon dioxide production (VCO2) are estimated. Outdoor air reference values are sampled after every 8 measurements. Gas sensors are calibrated prior to the onset of experiments with primary gas standards containing known concentrations of O2, CO2, and N2 (Airgas Puritan Medical, Ontario, CA). Respiratory exchange ratios (RER) were calculated as the ratio of carbon dioxide production (VCO2) to oxygen consumption (VO2). Energy expenditure measures (VO2, VCO2 and heat formation [(3.815 + 1.232*RER)*VO2 (in liters)]) were corrected for effective metabolic mass by using each mouse’s lean mass obtained from the EchoMRI test.

#### Open field test

This test predicts how animals respond when introduced into a brightly illuminated open arena ^141^. It is a classical test of "emotionality" used to measure anxiety-like responses of rodents exposed to stressful environmental stimuli (brightly illuminated open spaces) as well as to capture spontaneous activity measures. The apparatus is a square white Plexiglas (50 x 50 cm) open field illuminated to 600 lux in the center. Each animal is placed in the center of the field and several behavioral parameters (distance traveled, velocity, center time, frequency in center) are recorded during a 5-minute observation period and analyzed using Noldus Ethovision XT software.

#### Novel object recognition test

This test assays recognition memory while leaving the spatial location of the objects intact and is believed to involve the hippocampus, perirhinal cortex, and raphe nuclei ^142–144^. The basic principal is that animals explore novel environments and that with repeated exposure decreased exploration ensues (i.e., habituation; ^145^). A subsequent object substitution results in dishabituation of the previously habituated exploratory behavior (^145–147)^ and is expressed as a preferential exploration of the novel object relative to familiar features in the environment. Mice were individually habituated to a 51cm x 51cm x 39cm open field for 5 min. They were then be tested with two identical objects placed in the field for 5 min. After two such trials (each separated by 1 minute in a holding cage), the mouse was tested in the object novelty recognition test in which a novel object replaced one of the familiar objects. Behavior was video recorded and then scored for object contact time. The first time the mice were tested the objects used were clear plastic rectangular boxes filled with blue marbles and green plastic drink bottles filled with water and for the second test the objects were amber glass bottles and glass flasks filled with beige marbles. All objects were too tall for the mice to climb up on and too heavy for the mice to move.

#### Footprint Pattern Test

Basic gait measures can be assessed using simple footprint pattern analysis ^148, 149^. Non-toxic paint was applied to each mouse’s paws (a different color was used for front and back paws). The mouse was then placed at one end of a runway covered in paper and allowed to ambulate until their paws no longer left marks. Measurements were forelimb and hindlimb stride lengths (left and right) and front and back leg stride widths. Three full strides were averaged for each mouse’s values. Data were excluded from mice that did not make 3 measurable strides (i.e. they circled or stopped).

#### Barnes maze test

This is a spatial memory test ^150–152^ sensitive to impaired hippocampal function ^153^. Mice learn to find an escape chamber (19 x 8 x 7 cm) below one of twenty holes (5 cm diameter, 5 cm from perimeter) below an elevated brightly lit and noisy platform (75 cm diameter, elevated 58 cm above floor) using cues placed around the room. Spatial learning and memory were assessed across 4 trials (maximum time is 3 min) and then directly analyzed on the final (5^th^) probe trial in which the tunnel was removed and the time spent in each quadrant was determined, and the percent time spent in the target quadrant (the one originally containing the escape box) was compared with the average percent time in the other three quadrants. This is a direct test of spatial memory as there is no potential for local cues to be used in the mouse’s behavioral decision.

#### Grip strength test

Grip strength was measured with a mouse Grip Strength Meter (Columbus Instruments) according to the manufacturer’s instructions (User Manual 0167-007). All-limb measurements were performed with the angled grid attachment, pulling the mouse towards the meter by the tail after engagement of all limbs. Four consecutive measurements per mouse were taken and the highest three values were averaged, and data were expressed as newtons of peak force.

#### Hanging wire test

The hanging wire test allows for the assessment of grip strength and motor coordination ^154, 155^. Mice were held so that only their forelimbs contact an elevated metal bar (2 mm diameter, 45 cm long, 37 cm above the floor) held parallel to the table by a large ring stand and let go to hang. Each mouse was given three trials separated by 30 seconds. Each trial was scored as follows and the average for each mouse was calculated: 0 — fell off, 1 — hung onto the wire by two forepaws, 2 — hung onto the wire by two forepaws, but also attempted to climb onto the wire, 3 — hung onto the wire by two forepaws plus one or both hindpaws around the wire, 4 — hung onto the wire by all four paws plus tail wrapped, 5 — escaped (crawled to the ring stand and righted itself or climbed down the stand to the table). Latency to falling off was also measured up to a maximum of 30 s.

#### Rotarod test

Rotarod balancing requires a variety of proprioceptive, vestibular, and fine-tuned motor abilities as well as motor learning capabilities ^149^. An Accurotar rotarod apparatus (Omnitech Electronics, Inc., Columbus, OH) was used in these studies. A protocol was used whereby the rod starts in a stationary state and then begins to rotate with a constant acceleration of 10 rpm. When the mice were incapable of staying on the moving rod, they fell 38cm into a sanichip bedding filled chamber, breaking a photobeam in the process. The time of fall (translated to the speed at fall) was recorded by computer. The mice were tested in four sets of 3 trials, alternating directions between sets which were 30 min apart.

#### Treadmill test

The treadmill exhaustion test evaluates exercise capacity and endurance ^156^. Mice are motivated to run to exhaustion in order to escape a shock at the base of the treadmill. Mice were trained to run in three daily 5 min sessions in which stopping would result in the mice touching the back of the apparatus and experiencing a mild shock (200 msec pulses of electric current with pulse repetition rate of 3 times per second (3 Hz) and an intensity of 1 mA). The treadmill speed for training was 10 m/min (0.373 mph). For the exhaustion test, the speed was initially set at 10 m/min for 5 min, and was increased 2 m/min every 2 min up to a maximum speed of 46 m/min (1.7 mph). The mice were run until they were exhausted or the maximal speed was achieved (which would mean a maximum run time of 41 min). Exhaustion was defined as the inability of the animal to run on the treadmill for 10 sec despite receiving shocks, a maximum of 30 mild shocks. To prevent injury, the mice were monitored carefully and continually during each session, and immediately upon meeting the criterion for exhaustion the shock grid was turned off and the mouse was removed from the treadmill.

### Isolating Nuclei from Frozen Tissues

Flash-frozen in liquid nitrogen (stored at -80°C) is a common practice to preserve tissue and organs that are not to be processed immediately ^157–159^. Organs and tissues were collected from freshly dissected mice, snap frozen using liquid Nitrogen, and stored at – 80°C. Organs and tissues were then transferred to a pre-chilled mortar and laid on top of dry ice; liquid Nitrogen was poured over the frozen tissue and a pestle was used to grind and pulverized the sample until a uniformly fine powder was obtained. Pulverized sample was the aliquoted and returned to – 80°C. To extract nuclei from frozen aliquots, 500 ul homogenization buffer (Nuclear Isolation Buffer 1 (NIM1) consisting of 250 mM Sucrose, 25 mM KCL, 10 mM Tris-buffer pH 8.0, 5 mM MgCl2, 1 mM DTT, and 10% Triton X-100) were added to powdered tissue and transferred to the mixture a glass Dounce homogenizer and dounced ∼ 10 times (avoiding bubbles) on ice. Add homogenization buffer up to 1 mL and filter homogenization solution through a 40 mm cell strainer. Centrifuge filtered solution at 600xg (acceleration 4, deacceleration 4) for 4 min at 4° C. Aspirate supernatant and resuspend in 200 mL of PBS. Nuclei were then count on CellDrop FL (DeNovix) using 1:1 Acridine Orange /Propidium Iodide and homogenization solution. Samples were diluted in PBS to 1 million/mL to seed each well (∼30,000 cells in 30 mL/well) of 384 well plate (Perking Elmer PhenoPlate 384-well black, clear flat bottom Cat No. 6057300) pre-coated (1 mL/25 cm^2^) with poly-L-Lysine (50 mL/mg). Centrifuge plate at 4000xg for 15 min at 4° C and immediately added 60 mL of 4% PFA to each well and incubate for 15 min at 4° C. Followed by one wash of PBS then proceed with microscopic imaging of epigenetic landscapes.

### Isolating Peripheral Blood Mononuclear Cells (PBMC’s)

200 mL of blood was collected retroorbital for each mouse and immediately mixed with an equal volume of 50 mM EDTA (to prevent coagulation). Blood mixture was further diluted with 200 mL of PBS and carefully layering on top of 750 mL of Ficoll-Paque Plus (Millipore Sigma Cat. No. GE17-1440-02); followed by density gradient centrifugation at 700xg for 30 minutes at room temperature. The PBMC rich layer (cloudy phase) was carefully collected avoiding mixing the above and below layers and transferred to a new tube containing 10 mL of PBS. The PBMC mixture was centrifuged at 700xg for 20 minutes at room temperature and then removed supernatant and resuspend pellet in 1 mL of PBS. Isolated PBMC’s were counted manually using a hemocytometer (Hausser Scientific Cat. No. 3120). Samples were diluted in PBS to 1 million/mL to seed each well (∼30,000 cells in 30 mL/well) of 384 well plate (Perking Elmer PhenoPlate 384-well black, clear flat bottom Cat No. 6057300) pre-coated (1 mL/25 cm^2^) with poly-L-Lysine (50 mL/mg). Centrifuge plate at 4000xg for 15 min at 4° C and immediately added 60 mL of 4% PFA to each well and incubate for 15 min at 4° C followed by PBS wash.

### Doxorubicin treatment

Two months old C57BL/6 mice were intraperitoneal injected with doxorubicin (Santa Cruz Biotechnology, Cat. No. 25316-40-9) 10 mg/Kg. Doxorubicin was diluted in 150mM NaCl solution, and control mice were injected only with the vehicle solution (150mM NaCl). Fourteen days after treatment, mice were sacrificed and liver collected and immediately snap frozen in liquid nitrogen.

### Caloric restriction

Animal studies were conducted in accordance with approved protocols submitted through the respective Institutional Animal Care and Use Committees (IACUCs) at the University of Wisconsin Madison. Caloric Restriction mice: male C57BL/6N mice were individually housed under pathogen free conditions. Mice were randomized into control or restricted groups at 2 months of age and fed the AIN-93M semi-purified diet (BioServ) either a Control diet (95% ad libitum) or Restricted diet (25% less than control). The mice were sacrificed at 7 months of age and tissues were harvested, flash frozen and stored at -80°C.

### Liver cancers

Animal studies were conducted in accordance with approved protocols submitted through the respective Institutional Animal Care and Use Committees (IACUCs) at the University of California San Diego. C57BL/6(?) male mice were intraperitoneally injected with 25 mg/kg diethylnitrosamine (DEN; N0258, Sigma-Aldrich) at postnatal day 15. At 8 months of age the mice were then sacrificed and dissected tumor and non-tumor parts of the liver and samples were immediately fresh frozen using dry ice/2-methylbutane bath and then stored at – 80 °C. All tissue collection occurred during a 4-hour time frame (Zeitgeiber time 7-11; corresponding to 1pm-5pm) to minimize circadian effects on metabolism, proliferation, etc.

### Microscopic Imaging of Epigenetic Landscapes

Wells were blocked with 2% BSA in PBS 0.5% Triton X-100 for 1 hour at room temperature, and then incubated with primary antibody overnight at 4C and then washed with PBS 3x (5 minutes each at room temperature). Next, wells were incubated with secondary antibody overnight at 4C and then washed with PBS 3x (5 minutes each at room temperature). Antibodies were used at the following concentrations: Anti H3K27ac 1:1000 (Active Motif Cat No. 39685), and /or Anti H3K27me3 1:1000 (Active Motif Cat No. 39155), and/or Anti H3K4me1 1:1000 (Active Motif Cat No. 61633). Antibodies were detected using the following secondary antibodies for their appropriate hots: Goat anti-Rabbit IgG (H+L) Alexa Fluor™ 488 (Thermo Fisher Scientific Cat. No. A11034), and Donkey anti-Mouse IgG (H+L) Alexa Fluor™ 488 (Thermo Fisher Scientific Cat. No.

A31570). Wells were counterstained with DAPI (Thermo Fisher Scientific Cat. No. D1306) during the secondary antibody staining and plates were sealed with adhesive foil (VWR Cat. No. 60941-124). Cells/nuclei were imaged on either an Opera Phenix high-content screening system (PerkinElmer) or an IC200-KIC (Vala Sciences) using a 20x objective. At least five fields/well and a total of 9 z-stacks at a 1mm z-step for Opera Phenix and nine fields/well and a total of 10 z-stacks at a 1mm z-step for IC200 were acquired and five wells per mouse sample were imaged. Unless stated otherwise, at least three wells and a minimum of 300 cells for each condition were used for analysis.

### Image Feature Extraction

Image features were extracted for each single cell (nucleus) within a segmentation mask annotating cell nuclei on the images. The radial falloff ^160, 161^ artifact of the images was corrected using BaSiC Illumination correction on all channels as a preprocessing step^160^. The segmentation was performed using StarDist^[Citation error]^ applied to DAPI images. We defined cell nuclei as the segmentation masks after the removal of small objects with an area corresponding to a radius less than 3-7𝜇𝑚, assuming spherical shape, dependent on tissue and microscope used. The cell nuclei masks were applied to all channels to isolate raw images of each nucleus in each channel. Texture features (252 2-dimensional TAS features; 756 3D TAS features per epigenetic mark/channel) were calculated from each nucleus for each channel as previously described^60, 61^. In this study, 28 binarized images per epigenetic mark/channel were obtained by applying band-pass thresholding with the following intervals: (𝑣*_m_*, ∞), (𝑣*_m_* − 𝑣*_m_*𝑝, ∞), (𝑣*_m_* + 𝑣*_m_*𝑝, ∞), (𝑣*_m_* − 𝑣*_m_*𝑝, 𝑣*_m_* + 𝑣*_m_*𝑝), where 𝑣_"_ indicates mean pixel/voxel values within a cell nuclei mask, 𝑝 indicates a factor to determine the width of the band, ranging from 0.1 to 0.9 with the step of 0.1. One static interval and three width-variable intervals with nine factors derive 28 binarized images.

### ImAge Axis Construction

The ImAge axis was obtained using z-scored features of young and old groups based on their centroids or linear SVM models. The centroid-based axis was then defined by the vector from the centroid of the young group to the centroid of the old group. The SVM-based axis was defined by the vector orthogonal to the hyperplane optimized to separate young and old groups; the precise location of this axis can be defined as the intersection of the orthonormal vector to the hyperplane and the centroid of all data points. The calculation of centroids and optimization of SVM models were performed using all features (without feature selection) over all data points resampled using bootstrap (see the following section for details of bootstrapping). It is important to note that the ImAge axis does not deform the feature space in any way; it is a simple linear transposition and projection.

The robustness of the ImAge axis was validated in 100 iterations of tests, where 75% and 25% of data were randomly sampled without replacement for training (axis construction) and testing (projection onto the axis), respectively. This training step only involves the young and old groups. Importantly, the other age or condition groups (middle age, perturbations) were not included in the training.

At each iteration of training/testing, we resampled single cell data with bootstraps of 200 cells to obtain the average data point representative to each mouse sample with uncertainty of the average. The bootstrap was performed 1000 times for each mouse sample to equalize the number of data points per mouse. In our analysis, the utilization of bootstrap sampling is essential to examine the aging related changes in the cell composition. Throughout the aging process, the cell identity remains unchanged, while the composition of the cell distribution undergoes variations. For instance, individual T cells from a young and an older mouse may appear identical, but when sampling a bootstrap of 200 cells from blood samples, the composition of cells, or the ratio between different cell types, differs between young and old mice. Clustering the single cells do not reveal the difference between young and old samples but simply put the cells of the sample type together.

The ImAge orthogonal distance was defined by the distance to the ImAge axis. Thus, the orthogonal component can be considered the positive scalar value representing the distance of a datapoint to the ImAge axis. The orthogonal distance 𝑑_0_ for a data point can be given by 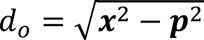, where 𝒙 is the relative coordinate of the datapoint with the origin of the ImAge axis, and 𝒑 is the coordinate of the same data point projected on the ImAge axis (i.e., 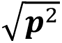 is the ImAge readout).

### Spearman correlation between multiple tissues

Spearman correlations were calculated via pairwise comparisons between all combinations of either 2 or 3 age groups per organ pair per channel. P-values were post-hoc corrected for multiple comparisons using the Bonferroni method. An alpha of 0.05 was applied to the p-values and significant correlations per organ pair per channel were counted. Organ pairs which were correlated in both all age groups and 2 age groups with the 3^rd^ removed were considered robust and significant.

### Information distance metric

We have used a distance metric based on mutual information. Considering two random variables 𝑥 and 𝑦 from corresponding distribution 𝑋 and 𝑌 that are normalized between 0 and 1 using min-max normalization, the mutual information 𝐼(𝑋, 𝑌) measures how much uncertainty of one variable is reduced by knowing the other variable. In other words, 𝐼(𝑋, 𝑌) measures the information shared by both variables. The variation of information 𝐷(𝑋, 𝑌) ^162^, or information distance metric we call here, is a measure of information that is not shared by both variables. Like the correlation and correlation distance, mutual information and information distance measures the dependence and independence between two random variables. However, unlike the correlation and correlation distance that measures the linear relationship between two variables, mutual information and information distance can capture the nonlinear relationship between two variables. The information distance can be written in terms of marginal entropies: 𝐻(𝑋), 𝐻(𝑌), and joint entropy 𝐻(𝑋, 𝑌)

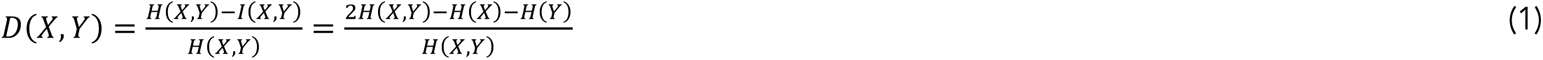

In practice, we can consider two samples as 𝑥 and 𝑦, with corresponding feature values 𝑥_!_ and 𝑦_!_ where 1 ≤ 𝑖 ≤ 𝑁*_feature_*. and 𝑁*_feature_*. is the number of features. The probability density 𝑋 and 𝑌, can be approximated by the histogram 𝑝(𝑧) of the feature values for each sample where 𝑝(𝑧) is the number density of feature values 𝑥*_i_* or 𝑦*_i_* that sit inside the bin centered at 𝑧 with bin size ℎ = 0.1. Entropy can be calculated using the Shannon entropy 𝐻(𝑧) = − ∑_z∈Z_ 𝑝(𝑧) logF𝑝(𝑧)G for 𝑧 = 𝑧_0_, 𝑧_1_.

### Embedding the data in the hyperbolic space

Hyperbolic geometry provides continuous approximation to tree-like hierarchical system, resulting in the distinguishing property of exponential expansion of states compared to quadratic expansion of states in Euclidean space ^163^. The hyperbolic geometry can be visualized using Poincare half space model and Poincare disk^163^. Hyperbolic space is better suited compared to Euclidean space for representing complex multipara metrical biological datasets. This is because hyperbolic space enables using fewer dimensions, inferring the hidden nodes based on the activity of observed “leaf” nodes, and reading out the “centrality” of each node ^164–166^. Using the information distance metric, the nodes in the middle are features that sample take different values and the nodes on the boundary are samples. In another words, the sample is a representation of a serious feature value selection starting from middle to boundary.

Considering the aging process is due to a sequence of biological changes, and each change is chosen from a variety of choices, hence the number of possible states increase exponentially. We employed the Hyperbolic Multidimensional Scaling (HMDS) ^67^, which is a version of MDS in hyperbolic space. Euclidean MDS minimizes the difference between the distances calculated in the original Euclidean data space and the distances calculated in the high dimensional reduced space. Once we specify the dimension of the reduced space, the quality of embedding could be assessed using Shepherd diagram, which compares the distance in the original space and the distance in the dimensionality reduced hyperbolic space.

In order to quantify the performance of the dimension reduction using MDS, we can plot the Shepherd diagram where we compare the distances between samples in the original high dimensional space and the distances between samples in the lower dimensional embedded space. In general, the better the MDS performs, the closer the distances between samples embedded in the lower dimension correlate linearly with the distances between samples in the original space. The linear relationship ensures that the distances between samples are not distorted when we embed the samples from original high dimensional space to lower dimensional space. The correlation, the tightness of linear fitting, ensures the accuracy and robustness of the embedding.

### Finding linear coefficients for behaviors

We took advantage of the logarithmic and exponential representations, which map the points from the hyperbolic space to the tangent space centered at a reference point and vice versa ^167^. The tangent space is Euclidean, and the curved geodesic in the hyperbolic space can be approximated by a straight line in the tangent space which is easier to compute.

In our analysis, we consider n samples with distinct biological ages represented as the column vector 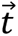 of dimension n. Similarly, we have N behavioral features, each denoted as 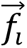, where 𝑖 ranges from 1 to N, also column vectors of dimension n. Instead of assessing the correlation between ImAge 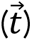 and individual readouts 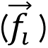, we explore the collective impact of these readouts on ImAge. We calculate the correlation 𝑅 between ImAge 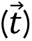 and the optimal linear combination of features, given by 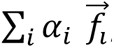, where {𝛼*_i_*} represents coefficients that maximize 𝑅, minimize p-values, and satisfy 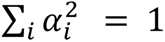. Non-significant and highly variable features have been systematically filtered, with a specific focus on those whose p-value, in terms of correlation with the ImAge variable, exceeds the threshold of 0.3. These features have been removed from the analysis, ensuring a more rigorous and refined dataset for subsequent investigations.

To ensure the uniqueness of the linear coefficients {𝛼*_i_*}, we compute a correlation matrix among all normalized significant feature vectors, grouping correlated feature vectors into orthogonal clusters. We identify 9 major orthogonal clusters of behavioral readouts, selecting a representative readout 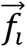, from each cluster for coefficient computation. To estimate feature variance and ensure consistency across multiple measurements, we applied four trials of jackknife resampling to each 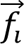, which encompasses several measurements over several days. Additionally, we conducted 1000 trials of reshuffling and recombination, such as selecting the first measurement from 𝑓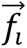 and the second measurement from 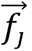. The linear coefficients {𝛼*_i_*} are computed as averages over these trials for each orthogonal cluster 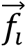.

### Separation between different cell types

To assess changes in epigenetic variation across diverse cell types during aging, we utilize two key metrics: the Silhouette score and the Kolmogorov–Smirnov distance (KS distance). The Silhouette score evaluates clustering performance, adaptable to both original and dimensional reduced data. In this study, we calculate Silhouette scores directly on raw data to avoid the bias introduced in the various dimension reduction processes. We utilize both the information distance and Euclidean distance metrics to illustrate changes in separation among different cell types from both an informational and a value perspective. Also, we employ 100 bootstraps to estimate the silhouette score within each age group, which allows us to assess the statistical stability and robustness of the silhouette score across various age categories.

To further confirm our observation of reduced separation between cell types during aging, we need a metric quantifying differences across cell types in individual features. The KS distance measures the distance between one-dimensional distributions characterizing distinct cell types. We sample 5000 cells per feature to capture distributions of specific cell types at given ages. Evaluating differences across 𝑁*_tissue_* tissue types follows a systematic approach: pairwise KS distances are computed between each tissue pair, with subsequent averaging. The resulting average KS distance is given by 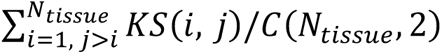 where 𝑖, 𝑗 denote cell types and the 𝐾𝑆(𝑖, 𝑗) is their KS distance and 𝐶(𝑁*_tissue_*, 2) is number of combinations choosing 2 pairwise tissues from total 𝑁*_tissue_* tissues

### Identification of signature cells

The signature cells for each age group (young or old) were identified as single cells where the ImAge readout was hardly observed in the other group and can be given by following indicator functions for young signature

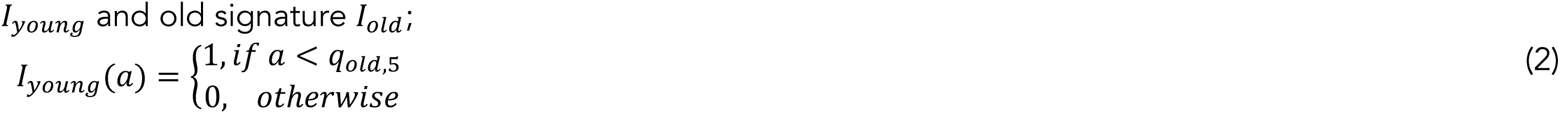

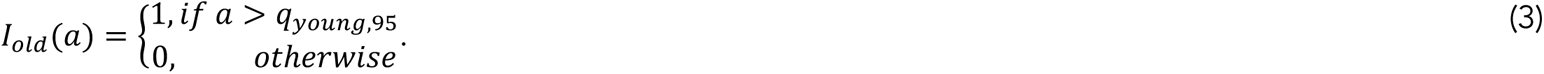

Here, 𝑎 indicates the ImAge readout for a single cell (nucleus), 𝑞*_old_*_,5_ indicates the 5^th^ percentile value of single-cell ImAge readout in old samples, and 𝑞*_young,95_* indicates the 95^th^ percentile value of single-cell ImAge readout in young samples. The proportion of signature cells and intermediate population can be obtained by calculating the proportion of them across all single-cell data obtained from each mouse sample. The choice of percentile values can adjust the purity of the definition of signature cells. For instance, using 0^th^ percentile (minimum value) of old in eq.(2) to define the young signature will ensure the selection of single cells only appear in the young group. We determined the young signatures by selecting the 5th percentile, which accounted for only 5 percent of young signature cells in the old group, considering these 5 percent as statistical noise. Similarly, we adopted the opposite approach to determine the old signature. This approach ensured that we had a sufficient number of signature cells without compromising the quality of the data by including too much statistical noise.

## ACKNOWLEDGEMENTS

We greatly appreciate the invaluable off Sanford Burnham Prebys animal and high throughput imaging core facilities personnel and leadership in particular Mary O’Rourke-Braxtan and Susanne Heynen-Genel (SBP NCI Cancer Center Support Grant P30 CA030199). We thank Dr. Collin Kaufman for his help with preparing tissues samples. We thank all members of the Terskikh laboratory for helpful discussion. This work was supported by NIH grants R21 AG068913 and R21 AG075483 to A.V.T.

## AUTHOR CONTRIBUTIONS

A.V.T., R.M.A., M.S, P.D.A., and T.O.S. designed and supervised the study; D.R., L.F., C.F., A.H, G.S.F., A.J.R., RMA and A.C. conducted the experiments; M.A.K., H.Q., K.N., W.M.H., D.R., T.O.S. and A.V.T. analyzed the data and prepared the manuscript.

## COMPETING FINANCIAL INTERESTS

The authors declare no competing financial interests.

## Extended Data Figure legends

**Extended Data Fig. 1:**
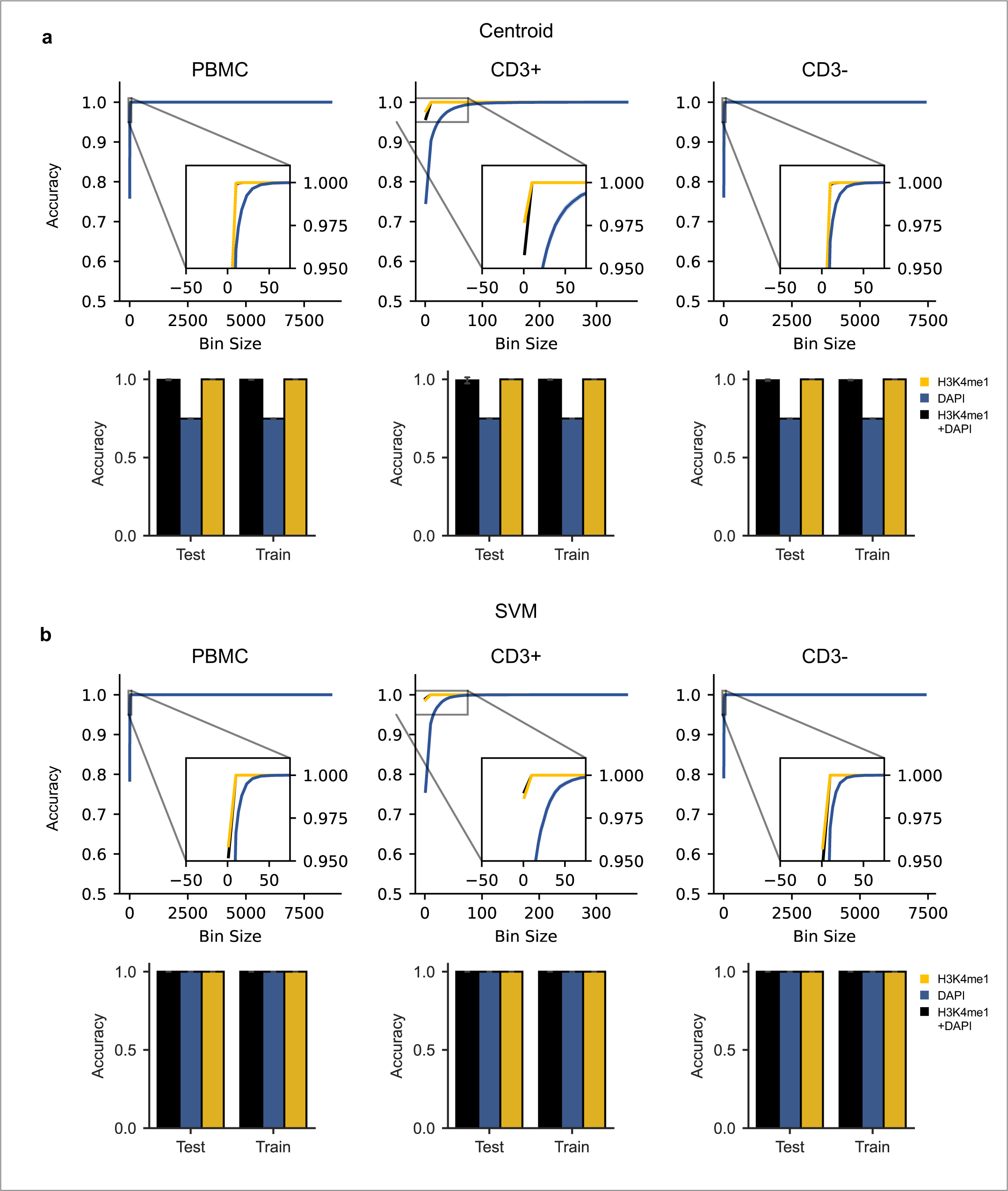
ImAge accuracy measurements in blood. **a-b** PBMCs were isolated from mice aged 2, 5, 9, 15, 21, 23, 32 months (n=2 for all groups), and immunolabeled for CD3 and H3K4me1 and costained with DAPI. **Lineplots:** Accuracy measurements versus bin size split by train and test data. 95% Confidence intervals are shown. **Barplots:** Final accuracy measurements for data shown in **figure 1** (200 cells per bin). Error bars represent +/- 2 sd. Measurements were made for all models constructed: **a**, centroid based **b**, SVM. PBMC: peripheral blood mononuclear cells. SVM: Support Vector Machine

**Extended Data Fig. 2:**
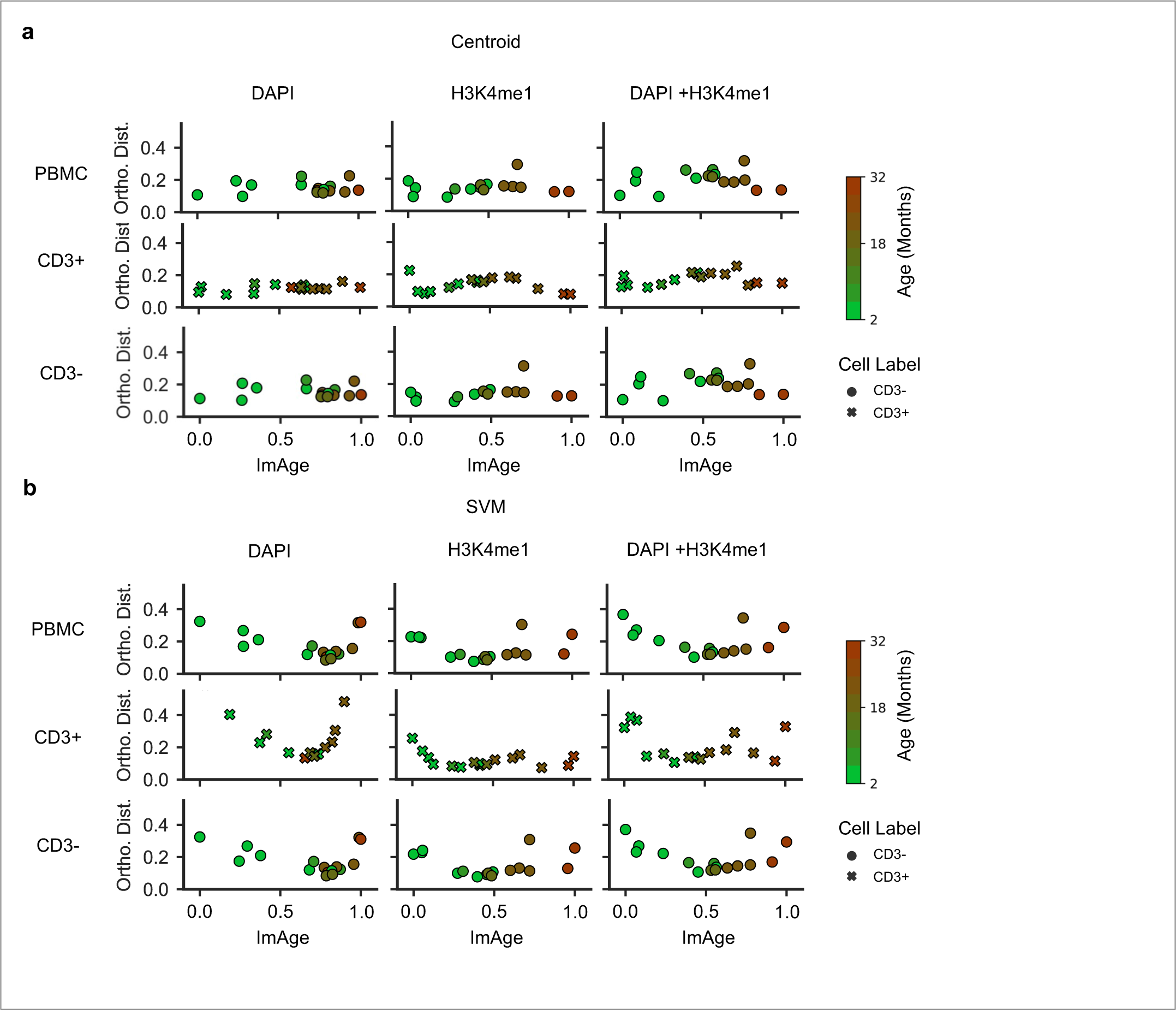
ImAge versus orthogonal distance plots. **a-b**, PBMCs were isolated from mice aged 2, 5, 9, 15, 21, 23, 32 months (n=2 for all groups), and immunolabeled for CD3 and H3K4me1 and costained with DAPI. Scatterplots show ImAge (normalized 0-1) versus orthogonal distance to the ImAge axis. ImAge was calculated using either **a**, centroid based method or **b**, SVM.

**Extended Data Fig. 3:**
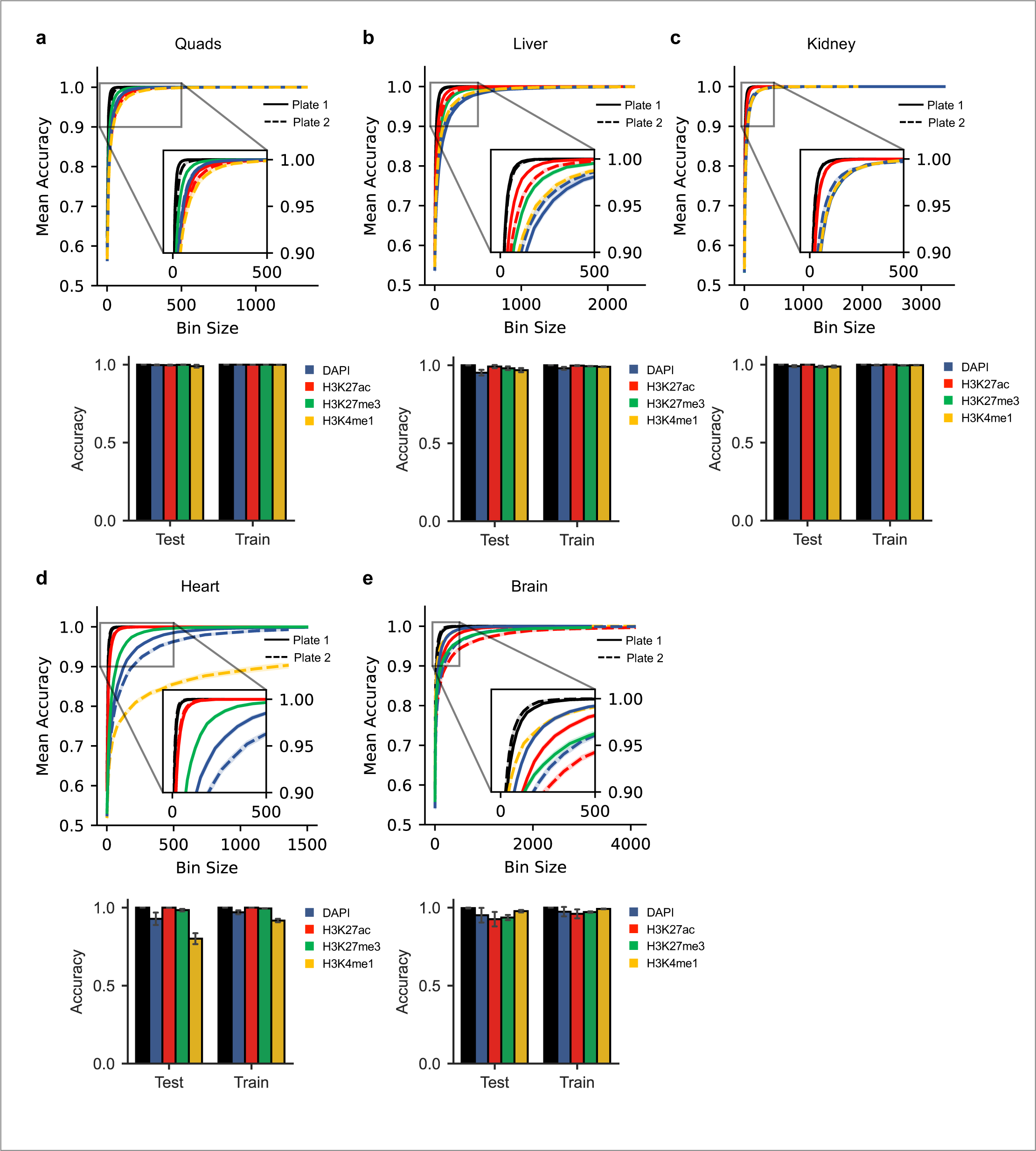
ImAge accuracy measurements in multiple solid organs. **A-E**, Nuclei isolated from solid organs of young (2 months, n=4) and aged (27 months, n=4) control mice were immunolabeled and imaged with two sets of antibodies: (H3K27ac & H3K4me1), (H3K27ac & H3K27me3), both costained with DAPI. **Lineplots**: Accuracy measurements versus bin size split by train and test data. 95% Confidence intervals are shown. Dashed or solid lines represent plate / antibody set of origin: (H3K27ac & H3K4me1), (H3K27ac & H3K27me3), respectively. **Barplots**: Final accuracy measurements for data shown in **figure 2** (200 cells per bin). Error bars represent +/- 2 sd. Measurements were made for all models constructed: **A**, quads **B**, liver **C**, kidney **D**, heart **E**, brain

**Extended Data Fig. 4:**
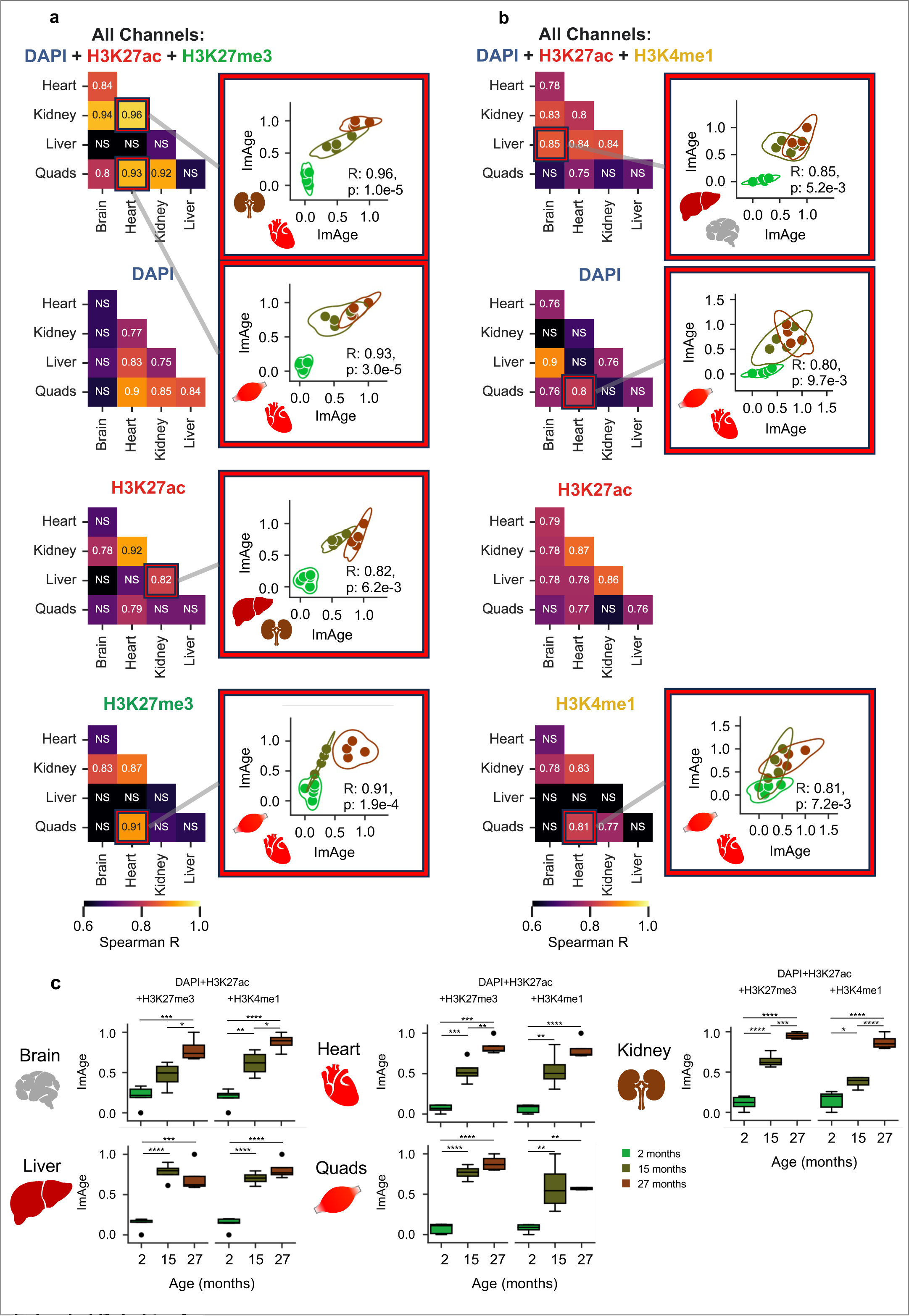
ImAge measures age in multiple solid organs. **A-B**, Pairwise spearman correlation coefficients of ImAge measurements from quadriceps (quads), liver, kidney, cardiac muscle (heart) and brain collected from three differentially aged cohorts of mice: 2 months (n=5) 15 months (n=4) and 27 months (n=4). Isolated nuclei were immunolabeled and imaged with two sets of antibodies, both costained with DAPI. The first antibody set visualized H3K27ac & H3K27me3 **(A)** and the second visualized H3K27ac & H3K4me1 **(B)**. Callouts show all organ pairs which were significantly correlated between all 3 age groups and at least 2 age groups leaving out the 3^rd^. KDE line in callouts represents the 5^th^ percentile. All ImAge data visualized is from the test split. P-values were Bonferroni corrected for multiple comparisons. KDE: Kernel Density Estimation

**Extended Data Fig. 5.**
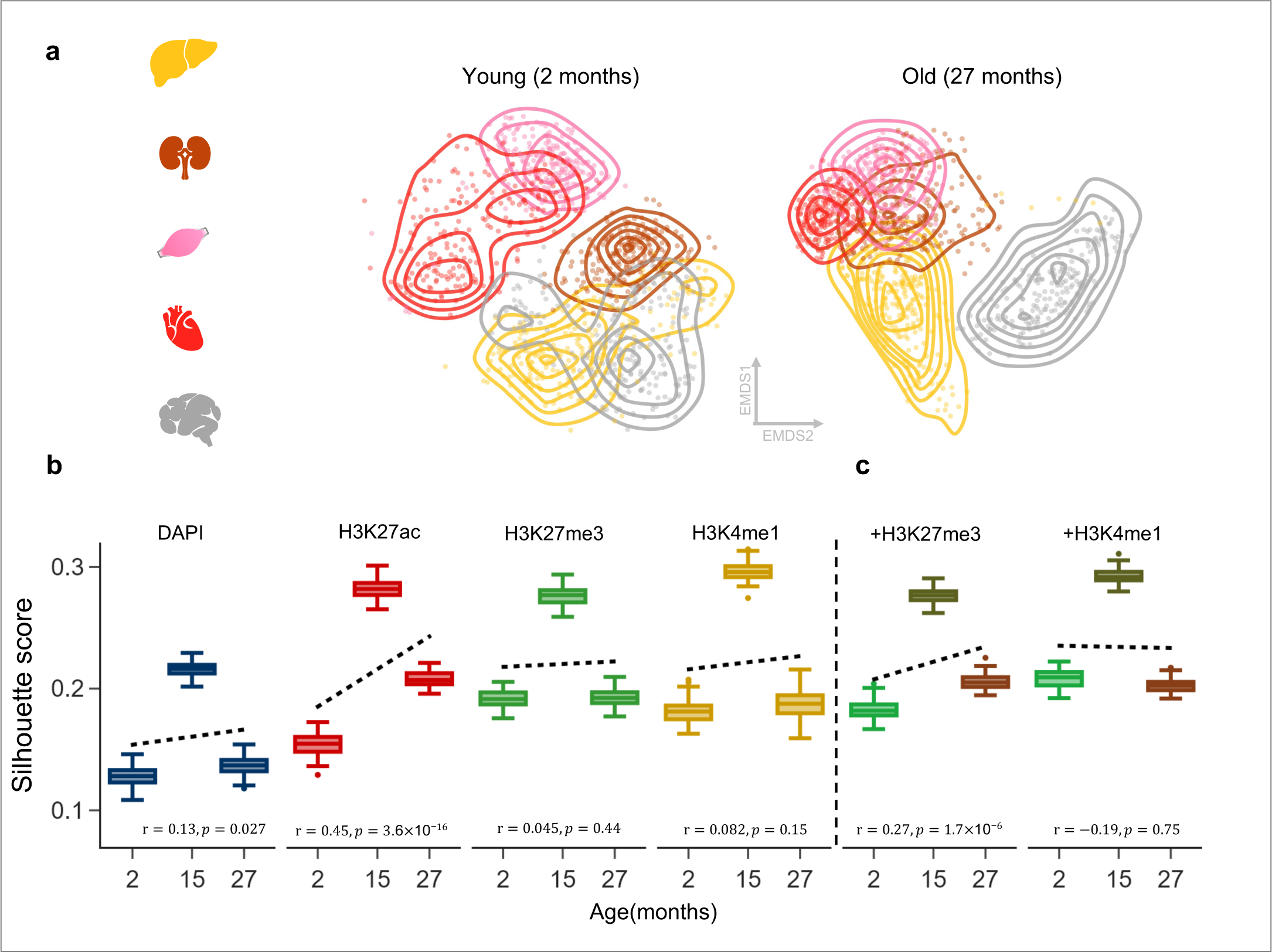
Separation between different cell types with aging including the Brain. **A** 2-dimensional EMDS of young (2 months) and old (27 months) liver, kidney, quads, and heart and brain. The observed clustering pattern reveals that the brain tissue cluster maintains a distinct separation from the clusters representing other tissue types, such as Kidney, Liver, Skeletal Muscle, and Cardiac Muscle. **B-C**. Silhouette scores of 5 organs at indicated ages for individual marks (b) or their combination (c) using the information distance metric. The Silhouette scores do not indicate a decline in tissue differentiation with aging in the presence of brain tissue due to slower progression of chromatin aging.

**Extended Data Fig. 6.**
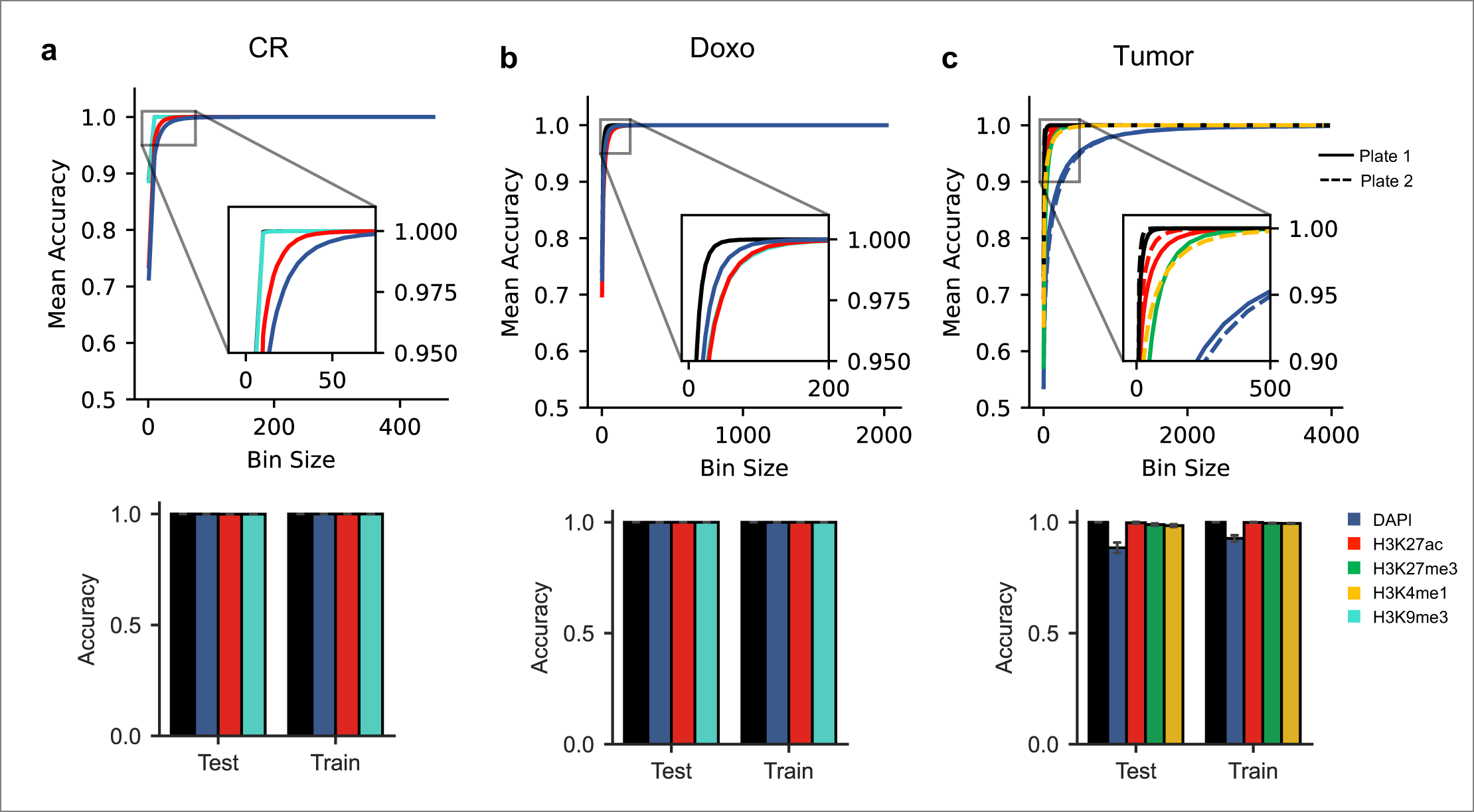
ImAge accuracy metrics for interventions in biological age. **A-C**, Nuclei from young and aged control mice were immunolabeled for H3K27ac and either H3K9me3 **(A & B)** or H3K27me3 & H3k4me1 **(C)** and costained with DAPI. **Lineplots:** Accuracy measurements versus bin size split by train and test data. 95% Confidence intervals are shown. **Barplots:** Final accuracy measurements for data shown in figure 4 (200 cells per bin). Error bars represent +/- 2 sd. **A**, Accuracy measurements for separating 3 month (n=3) and 24 month (n=3) old mice at various bin sizes. **B**, Accuracy measurements for separating 1 month (n=4) and 27 month (n=4) old mice at various bin sizes. **C**, Accuracy measurements for separating 2 month (n=5) and 8 month (n=6) old mice at various bin sizes. Dashed or solid lines represent plate / antibody set of origin: (H3K27ac & H3K4me1), (H3K27ac & H3K27me3).

**Extended Data Fig. 7.**
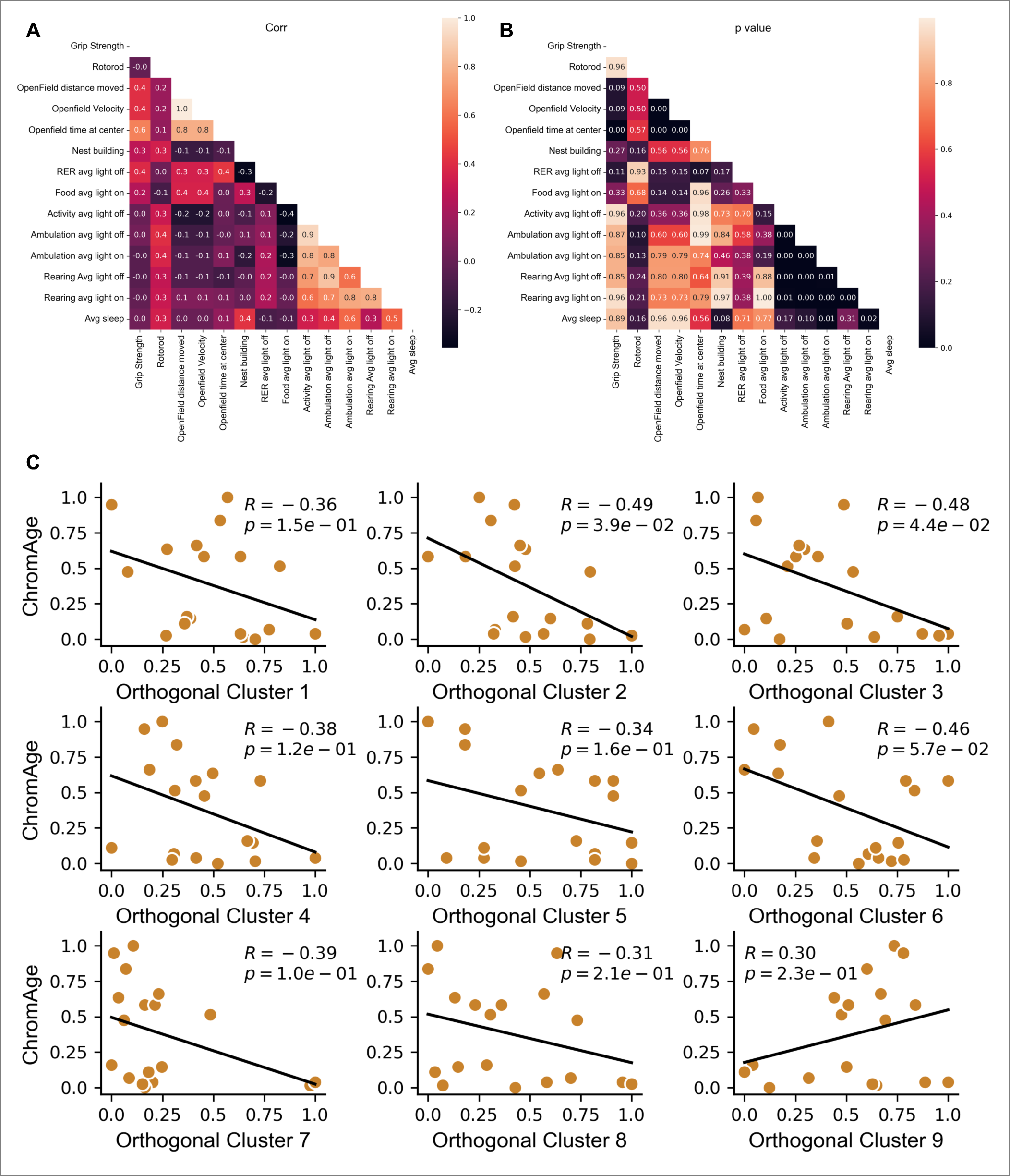
A-B Clustering of individual features and individual behavior clusters correlations. To make our linear coefficients 𝜶_𝒊_ unique, we need to find orthogonal bases from individual behaviors. The criterion for behaviors clustering together is based on the **A** high Pearson correlation (>0.7) and high significance according to **B** p values (<0.05).**C** Correlation between ImAge and clusters of behaviors. Investigating the relationship between ImAge and individual orthogonal clusters of behaviors.

**Extended Data Fig. 8.**
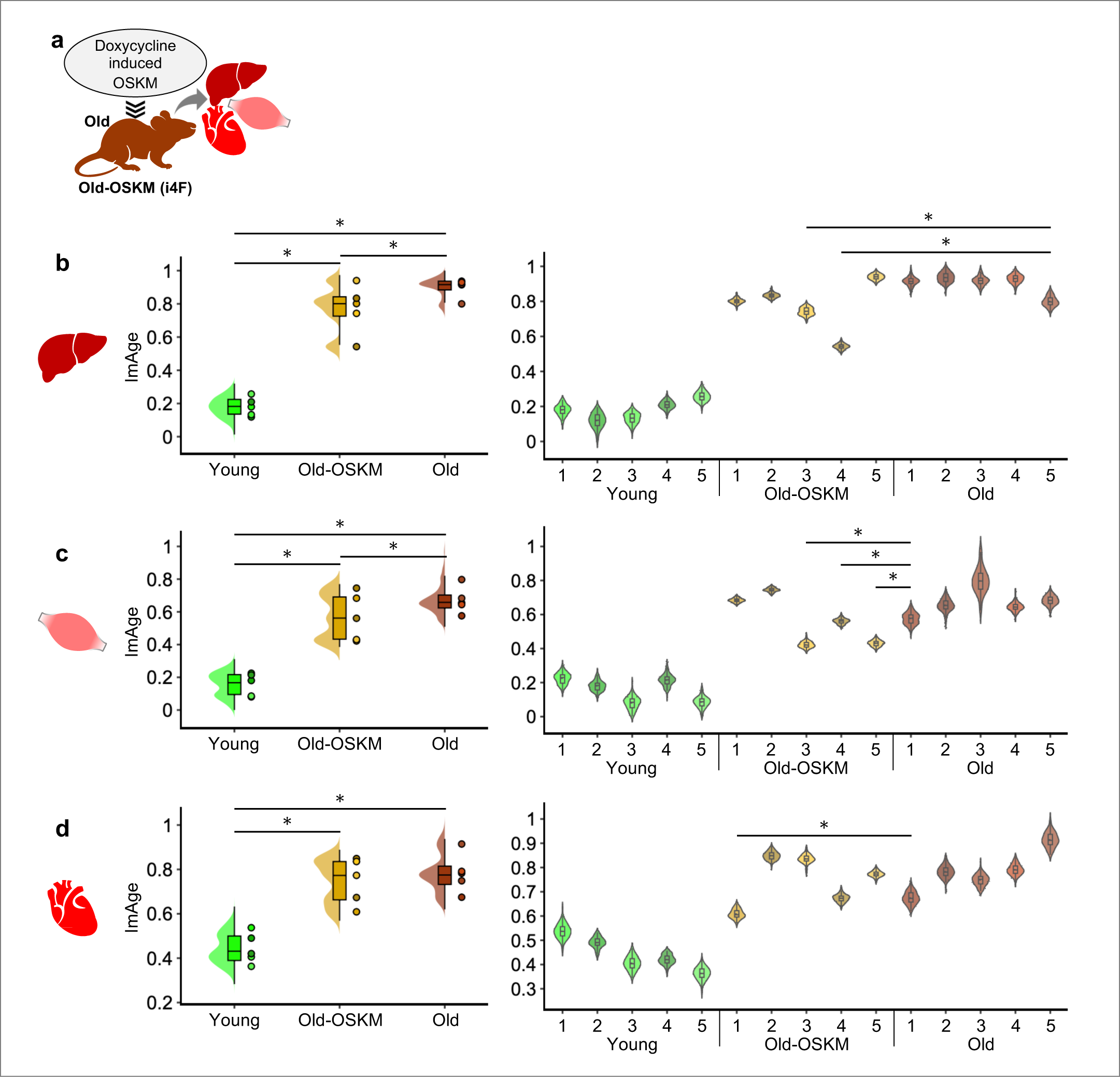
ImAge detected partial reprogramming of multiple organs at mouse sample level and enabled to find signatures of different age groups. The chronological ages of young and old mice are 3.2 and 13.8 months, respectively. **a**, A graphical representation of the experimental condition; old mice were treated with doxycycline to overexpress OSKM factors to evaluate the degree of reprogramming on the liver, quadriceps and cardiac muscle (heart). **b-d (left)**, Distribution of ImAge obtained from 100 iterations of the test procedure using the linear support vector machine. Round-shaped symbols on the right side of violin plots for each age group represent the mean ImAge of each mouse sample. **b-d (right)**, Sample-wise comparison of ImAge. Each violin plot represents the distribution of the ImAge for each mouse sample with the numbers from one to five for young, Old-OSKM and old groups. In all panels, an asterisk indicates the significant decrease from the old mouse showed the lowest ImAge in the old group with respect to median ImAge (Mann-Whitney U-test, significant threshold; p<0.05).

**Extended Data Fig. 9.**
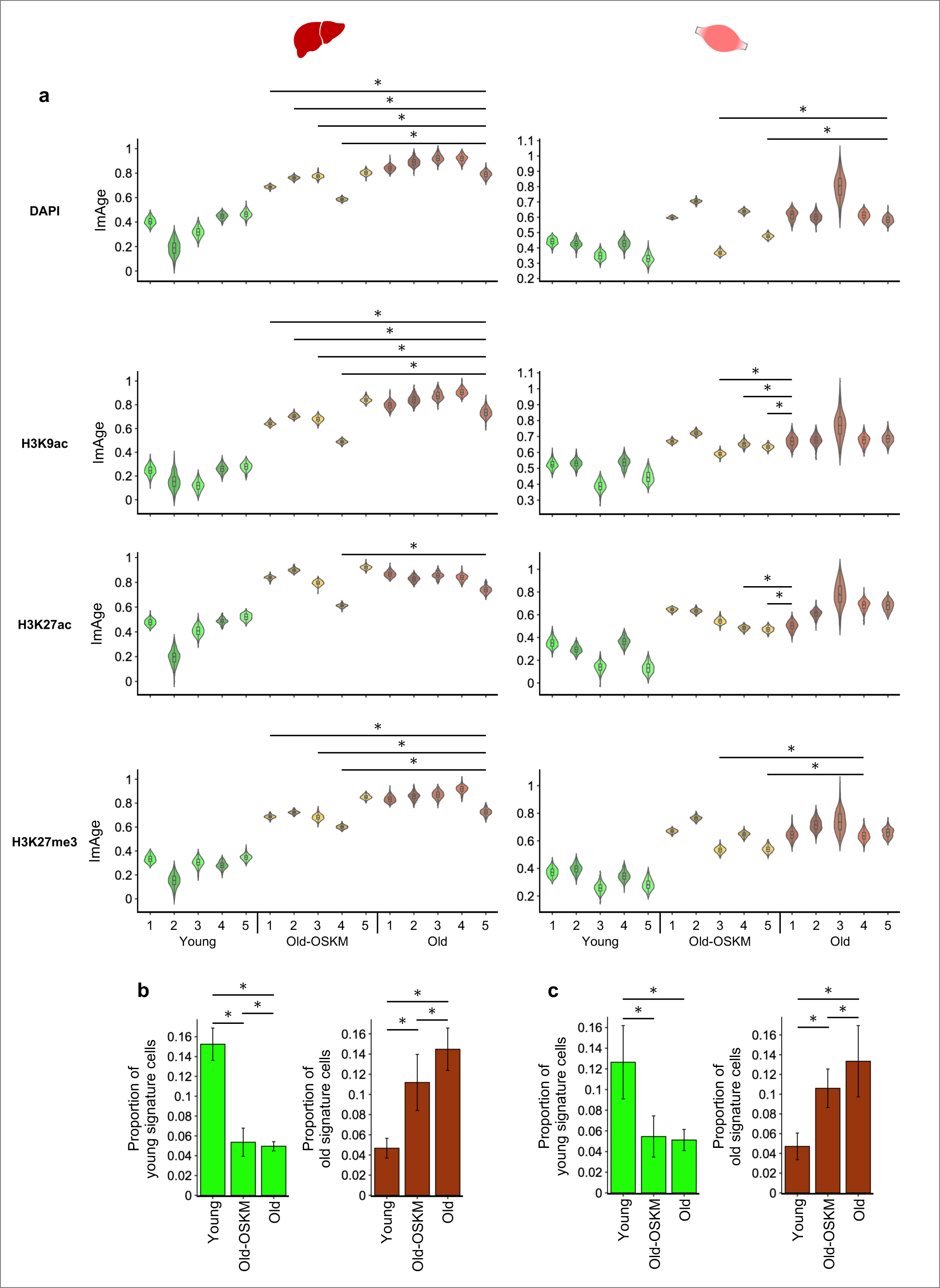
Degree of the partial reprogramming varies depending on tissues and epigenetic marks/channels. The chronological ages of young and old mice are 3.2 and 13.8 months, respectively. **a**, Each violin plot represents the distribution of the ImAge for each mouse sample with the numbers from one to five for young, Old-OSKM and old groups. Left and right columns are for liver and muscle, respectively. In all panels, an asterisk indicates the significant decrease from the old mouse showed the lowest ImAge in the old group with respect to median ImAge (Mann-Whitney U-test, p<0.05). **b, c**, proportion of cells with young and old signature ImAge in each age group in the liver (b) and muscle (c) (Mann-Whitney U-test, p<0.05).

**Extended Data Fig. 10.**
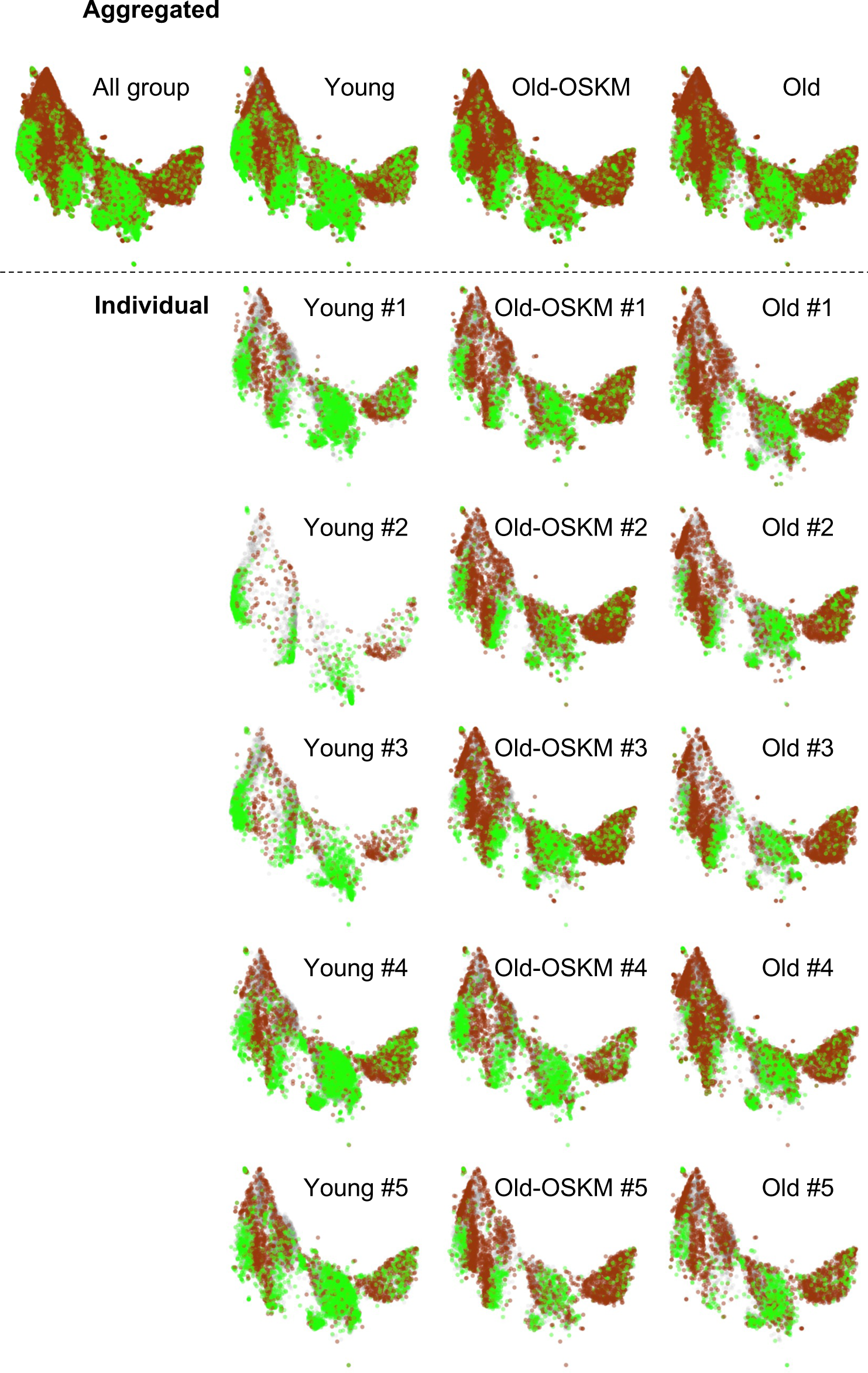
Overview of young/old signature of ImAge revealed at the single cell level in liver samples with in-vivo partial reprogramming. uniform manifold approximation and projection (UMAP) of three-dimensional threshold threshold adjacency statistic (TAS) features for single cells. The green and brown data points represent the single cells with signature ImAge of young and old, respectively. The gray points represent the intermediate single cells that did not belongs to either of the signatures.

## Notes

### Competing Interest Statement

The authors have declared no competing interest.

### Summary of Updates

This is the major revisions to include the analysis of in vivo reprogramming and to update previous findings

## REFERENCES

1. Zampino, M. et al. Biomarkers of aging in real life: three questions on aging and the comprehensive geriatric assessment. GeroScience 44, 2611–2622 (2022).

2. Baker, G. T. & Sprott, R. L. Biomarkers of aging. Experimental Gerontology 23, 223–239 (1988).

3. Wagner, K.-H., Cameron-Smith, D., Wessner, B. & Franzke, B. Biomarkers of aging: from function to molecular biology. Nutrients 8, 338 (2016).

4. LaPierre, T. A. & Hughes, M. E. Population aging in Canada and the United States. in International handbook of population aging 191–230 (Springer, 2009).

5. Control, Centers for Disease & Prevention. Trends in aging--United States and worldwide. MMWR. Morbidity and mortality weekly report 52, 101–106 (2003).

6. Jylhävä, J., Pedersen, N. L. & Hägg, S. Biological Age Predictors. EBioMedicine 21, 29–36 (2017).

7. Jones, J. A. B. et al. The AgeGuess database, an open online resource on chronological and perceived ages of people aged 5–100. Scientific Data 6, 1–8 (2019).

8. Simm, A. et al. Potential biomarkers of ageing. Biol. Chem. 389, 257–265 (2008).

9. Jazwinski, S. M. & Kim, S. Examination of the Dimensions of Biological Age. Front. Genet. 10, 263 (2019).

10. Rantanen, T. et al. Muscle strength and body mass index as long-term predictors of mortality in initially healthy men. The Journals of Gerontology Series A: Biological Sciences and Medical Sciences 55, M168– M173 (2000).

11. Whitehead, J. C. et al. A clinical frailty index in aging mice: comparisons with frailty index data in humans. Journals of Gerontology Series A: Biomedical Sciences and Medical Sciences 69, 621–632 (2014).

12. Kane, A. E., Keller, K. M., Heinze-Milne, S., Grandy, S. A. & Howlett, S. E. A murine frailty index based on clinical and laboratory measurements: links between frailty and pro-inflammatory cytokines differ in a sex-specific manner. The Journals of Gerontology: Series A 74, 275–282 (2019).

13. Schultz, M. B. et al. Age and life expectancy clocks based on machine learning analysis of mouse frailty. Nat. Commun. 11, 4618 (2020).

14. Alpert, A. et al. A clinically meaningful metric of immune age derived from high-dimensional longitudinal monitoring. Nature medicine 25, 487–495 (2019).

15. de Toda, I. M., Vida, C., San Miguel, L. S. & De la Fuente, M. When will my mouse die? Life span prediction based on immune function, redox and behavioural parameters in female mice at the adult age. Mechanisms of Ageing and Development 182, 111125 (2019).

16. Mather, K. A., Jorm, A. F., Parslow, R. A. & Christensen, H. Is telomere length a biomarker of aging? A review. Journals of Gerontology Series A: Biomedical Sciences and Medical Sciences 66, 202–213 (2011).

17. Krištić, J. et al. Glycans are a novel biomarker of chronological and biological ages. Journals of Gerontology Series A: Biomedical Sciences and Medical Sciences 69, 779–789 (2014).

18. Wang, A. S. & Dreesen, O. Biomarkers of cellular senescence and skin aging. Frontiers in Genetics 9, 247 (2018).

19. Hannum, G. et al. Genome-wide methylation profiles reveal quantitative views of human aging rates. Mol. Cell 49, 359–367 (2013).

20. Horvath, S. DNA methylation age of human tissues and cell types. Genome Biol. 14, R115 (2013).

21. Lowe, R. et al. Ageing-associated DNA methylation dynamics are a molecular readout of lifespan variation among mammalian species. Genome Biol. 19, 22 (2018).

22. Galow, A.-M. & Peleg, S. How to Slow down the Ticking Clock: Age-Associated Epigenetic Alterations and Related Interventions to Extend Life Span. Cells 11, (2022).

23. Lu, A. T. et al. DNA methylation GrimAge strongly predicts lifespan and healthspan. Aging 11, 303–327 (2019).

24. Weidner, C. I. et al. Aging of blood can be tracked by DNA methylation changes at just three CpG sites. Genome Biol. 15, R24 (2014).

25. Levine, M. E. et al. An epigenetic biomarker of aging for lifespan and healthspan. Aging 10, 573–591 (2018).

26. Bell, C. G. et al. DNA methylation aging clocks: challenges and recommendations. Genome Biol. 20, 249 (2019).

27. Lu, A. T., et al. Universal DNA methylation age across mammalian tissues. *bioRxiv* 2021.01.18.426733 (2022) doi:10.1101/2021.01.18.426733.

28. Levine, M. E., Higgins-Chen, A., Thrush, K., Minteer, C. & Niimi, P. Clock work: Deconstructing the epigenetic clock signals in aging, disease, and reprogramming. bioRxiv (2022) doi:10.1101/2022.02.13.480245.

29. Liu, Z. et al. Underlying features of epigenetic aging clocks in vivo and in vitro. Aging Cell 19, e13229 (2020).

30. Field, A. E. et al. DNA Methylation Clocks in Aging: Categories, Causes, and Consequences. Mol. Cell 71, 882–895 (2018).

31. Feridooni, H. A. et al. The impact of age and frailty on ventricular structure and function in C57BL/6J mice. The Journal of physiology 595, 3721–3742 (2017).

32. Rockwood, K. et al. A frailty index based on deficit accumulation quantifies mortality risk in humans and in mice. Scientific reports 7, 43068 (2017).

33. Kane, A. E. et al. Impact of longevity interventions on a validated mouse clinical frailty index. Journals of Gerontology Series A: Biomedical Sciences and Medical Sciences 71, 333–339 (2016).

34. Wang, T. et al. Epigenetic aging signatures in mice livers are slowed by dwarfism, calorie restriction and rapamycin treatment. Genome Biol. 18, 57 (2017).

35. Williams, T. D. Individual variation in endocrine systems: moving beyond the “tyranny of the Golden Mean.” Philos. Trans. R. Soc. Lond. B Biol. Sci. 363, 1687–1698 (2008).

36. Amundson, R. Against normal function. Studies in History and Philosophy of Science Part C: Studies in History and Philosophy of Biological and Biomedical Sciences 31, 33–53 (2000).

37. Westneat, D. F., Wright, J. & Dingemanse, N. J. The biology hidden inside residual within-individual phenotypic variation. Biol. Rev. Camb. Philos. Soc. 90, 729–743 (2015).

38. Zhavoronkov, A. et al. Artificial intelligence for aging and longevity research: Recent advances and perspectives. Ageing research reviews 49, 49–66 (2019).

39. Yang, J.-H. et al. Loss of Epigenetic Information as a Cause of Mammalian Aging. Cell (2023) doi:10.1016/j.cell.2022.12.027.

40. Takahashi, K. & Yamanaka, S. Induction of pluripotent stem cells from mouse embryonic and adult fibroblast cultures by defined factors. Cell 126, 663–676 (2006).

41. Ocampo, A. et al. In Vivo Amelioration of Age-Associated Hallmarks by Partial Reprogramming. Cell vol. 167 1719–1733.e12 (2016).

42. Chondronasiou, D. et al. Multi-omic rejuvenation of naturally aged tissues by a single cycle of transient reprogramming. Aging Cell 21, e13578 (2022).

43. Ng, R. K. & Gurdon, J. B. Epigenetic inheritance of cell differentiation status. Cell Cycle 7, 1173–1177 (2008).

44. Barrero, M. J., Boué, S. & Izpisúa Belmonte, J. C. Epigenetic mechanisms that regulate cell identity. Cell Stem Cell 7, 565–570 (2010).

45. Ma, S. et al. Chromatin Potential Identified by Shared Single-Cell Profiling of RNA and Chromatin. Cell 183, 1103–1116.e20 (2020).

46. Adam, R. C. et al. Temporal Layering of Signaling Effectors Drives Chromatin Remodeling during Hair Follicle Stem Cell Lineage Progression. Cell Stem Cell 22, 398–413.e7 (2018).

47. Song, M.-R. & Ghosh, A. FGF2-induced chromatin remodeling regulates CNTF-mediated gene expression and astrocyte differentiation. Nat. Neurosci. 7, 229–235 (2004).

48. Meshorer, E. Chromatin in embryonic stem cell neuronal differentiation. Histol. Histopathol. 22, 311–319 (2007).

49. Arnsdorf, E. J., Tummala, P., Castillo, A. B., Zhang, F. & Jacobs, C. R. The epigenetic mechanism of mechanically induced osteogenic differentiation. J. Biomech. 43, 2881–2886 (2010).

50. Morales Berstein, F., et al. Assessing the causal role of epigenetic clocks in the development of multiple cancers: a Mendelian randomization study. Elife 11, (2022).

51. Tsurumi, A. & Li, W. X. Global heterochromatin loss: a unifying theory of aging? Epigenetics 7, 680–688 (2012).

52. Lee, J.-H., Kim, E. W., Croteau, D. L. & Bohr, V. A. Heterochromatin: an epigenetic point of view in aging. Exp. Mol. Med. 52, 1466–1474 (2020).

53. Villeponteau, B. The heterochromatin loss model of aging. Exp. Gerontol. 32, 383–394 (1997).

54. Sen, P., Shah, P. P., Nativio, R. & Berger, S. L. Epigenetic Mechanisms of Longevity and Aging. Cell 166, 822–839 (2016).

55. Creyghton, M. P. et al. Histone H3K27ac separates active from poised enhancers and predicts developmental state. Proc Natl Acad Sci U S A 107, 21931–21936 (2010).

56. Ucar, D. et al. The chromatin accessibility signature of human immune aging stems from CD8(+) T cells. J Exp Med 214, 3123–3144 (2017).

57. Maybury-Lewis, S. Y. et al. Changing and stable chromatin accessibility supports transcriptional overhaul during neural stem cell activation and is altered with age. Aging Cell 20, e13499 (2021).

58. Farhy, C. et al. Improving drug discovery using image-based multiparametric analysis of the epigenetic landscape. Elife 8, (2019).

59. Weigert, M., Schmidt, U., Haase, R., Sugawara, K. & Myers, G. Star-convex polyhedra for 3d object detection and segmentation in microscopy. in Proceedings of the IEEE/CVF Winter Conference on Applications of Computer Vision 3666–3673 (2020).

60. Hamilton, N. A., Pantelic, R. S., Hanson, K. & Teasdale, R. D. Fast automated cell phenotype image classification. BMC Bioinformatics 8, 110 (2007).

61. Tahir, M., Jan, B., Hayat, M., Shah, S. U. & Amin, M. Efficient computational model for classification of protein localization images using Extended Threshold Adjacency Statistics and Support Vector Machines. Comput. Methods Programs Biomed. 157, 205–215 (2018).

62. Becht, E. et al. Dimensionality reduction for visualizing single-cell data using UMAP. Nat. Biotechnol. (2018) doi:10.1038/nbt.4314.

63. Zhou, Y. & Sharpee, T. O. Hyperbolic geometry of gene expression. iScience 24, 102225 (2021).

64. Praturu, A. & Sharpee, T. A Bayesian approach to hyperbolic Multi-Dimensional Scaling. bioRxiv (2022) doi:10.1101/2022.10.12.511940.

65. Van der Maaten, L. & Hinton, G. Visualizing data using t-SNE. J. Mach. Learn. Res. 9, (2008).

66. Zhou, Y., Smith, B. H. & Sharpee, T. O. Hyperbolic geometry of the olfactory space. Science Advances 4, eaaq1458 (2018).

67. Praturu, A. & Sharpee, T. A Bayesian Approach to Hyperbolic Embeddings. Bull. Am. Phys. Soc. (2022).

68. Zhang, H., Rich, P. D., Lee, A. K. & Sharpee, T. O. Hippocampal spatial representations exhibit a hyperbolic geometry that expands with experience. Nat. Neurosci. 26, 131–139 (2023).

69. Zhou, Y. & Sharpee, T. O. Using global t-SNE to preserve inter-cluster data structure. bioarxiv (2018).

70. Piening, B. D., Lovejoy, J. & Earls, J. C. Ageotypes: Distinct Biomolecular Trajectories in Human Aging. Trends in pharmacological sciences vol. 41 299–301 (2020).

71. Ahadi, S. et al. Personal aging markers and ageotypes revealed by deep longitudinal profiling. Nat. Med. 26, 83–90 (2020).

72. Baar, M. P. et al. Targeted Apoptosis of Senescent Cells Restores Tissue Homeostasis in Response to Chemotoxicity and Aging. Cell vol. 169 132–147.e16 (2017).

73. Cupit-Link, M. C. et al. Biology of premature ageing in survivors of cancer. ESMO Open 2, e000250 (2017).

74. Hill, A., Sadda, J., LaBarge, M. A. & Hurria, A. How cancer therapeutics cause accelerated aging: Insights from the hallmarks of aging. J. Geriatr. Oncol. 11, 191–193 (2020).

75. Demaria, M. et al. Cellular Senescence Promotes Adverse Effects of Chemotherapy and Cancer Relapse. Cancer Discov. 7, 165–176 (2017).

76. Severgnini, M. et al. A rapid two-step method for isolation of functional primary mouse hepatocytes: cell characterization and asialoglycoprotein receptor based assay development. Cytotechnology 64, 187–195 (2012).

77. Madeo, F., Carmona-Gutierrez, D., Hofer, S. J. & Kroemer, G. Caloric Restriction Mimetics against Age-Associated Disease: Targets, Mechanisms, and Therapeutic Potential. Cell Metab. 29, 592–610 (2019).

78. Ma, S. & Gladyshev, V. N. Molecular signatures of longevity: Insights from cross-species comparative studies. Semin. Cell Dev. Biol. 70, 190–203 (2017).

79. Balasubramanian, P., Howell, P. R. & Anderson, R. M. Aging and Caloric Restriction Research: A Biological Perspective With Translational Potential. EBioMedicine 21, 37–44 (2017).

80. Lacar, B. et al. Nuclear RNA-seq of single neurons reveals molecular signatures of activation. Nat. Commun. 7, 11022 (2016).

81. Tomimatsu, K. et al. Locus-specific induction of gene expression from heterochromatin loci during cellular senescence. Nat Aging 2, 31–45 (2022).

82. Levine, M. et al. A rat epigenetic clock recapitulates phenotypic aging and co-localizes with heterochromatin. Elife 9, (2020).

83. Wang, K. et al. Epigenetic regulation of aging: implications for interventions of aging and diseases. Signal Transduct Target Ther 7, 374 (2022).

84. Baskin, K. K., Winders, B. R. & Olson, E. N. Muscle as a “mediator” of systemic metabolism. Cell metabolism 21, 237–248 (2015).

85. Puri, D. & Wagner, W. Epigenetic rejuvenation by partial reprogramming. Bioessays 45, e2200208 (2023).

86. Horvath, S. & Raj, K. DNA methylation-based biomarkers and the epigenetic clock theory of ageing. Nat. Rev. Genet. 19, 371–384 (2018).

87. Finn, E. H. & Misteli, T. Molecular basis and biological function of variability in spatial genome organization. Science 365, (2019).

88. Trapp, A., Kerepesi, C. & Gladyshev, V. N. Profiling epigenetic age in single cells. Nat Aging 1, 1189– 1201 (2021).

89. Schaum, N. et al. Ageing hallmarks exhibit organ-specific temporal signatures. Nature 583, 596–602 (2020).

90. Nie, C. et al. Distinct biological ages of organs and systems identified from a multi-omics study. Cell Rep. 38, 110459 (2022).

91. Nishi, H., Higashihara, T. & Inagi, R. Lipotoxicity in Kidney, Heart, and Skeletal Muscle Dysfunction. Nutrients 11, (2019).

92. Slawik, M. & Vidal-Puig, A. J. Lipotoxicity, overnutrition and energy metabolism in aging. Ageing Res. Rev. 5, 144–164 (2006).

93. Samarghandian, S., Azimi-Nezhad, M., Farkhondeh, T. & Samini, F. Anti-oxidative effects of curcumin on immobilization-induced oxidative stress in rat brain, liver and kidney. Biomed. Pharmacother. 87, 223– 229 (2017).

94. Abarikwu, S. O. Protective effect of quercetin on atrazine-induced oxidative stress in the liver, kidney, brain, and heart of adult wistar rats. Toxicol. Int. 21, 148–155 (2014).

95. Bahar, R. et al. Increased cell-to-cell variation in gene expression in ageing mouse heart. Nature 441, 1011–1014 (2006).

96. Tauc, H. M. et al. Age-related changes in polycomb gene regulation disrupt lineage fidelity in intestinal stem cells. Elife 10, (2021).

97. Stewart, T. M., Holbert, C. E. & Casero, R. A., Jr. Helping the helpers: polyamines help maintain helper T-cell lineage fidelity. Immunometabolism (Cobham) 4, e00002 (2022).

98. Liu, Z. et al. Ectopic resurrection of embryonic/developmental genes in aging. Current Medicine 1, 11 (2022).

99. Vergnes, L., Péterfy, M., Bergo, M. O., Young, S. G. & Reue, K. Lamin B1 is required for mouse development and nuclear integrity. Proc. Natl. Acad. Sci. U. S. A. 101, 10428–10433 (2004).

100. Liu, X. et al. Resurrection of endogenous retroviruses during aging reinforces senescence. Cell (2023) doi:10.1016/j.cell.2022.12.017.

101. Simon, M. et al. LINE1 Derepression in Aged Wild-Type and SIRT6-Deficient Mice Drives Inflammation. Cell Metab. 29, 871–885.e5 (2019).

102. Oberdoerffer, P. & Sinclair, D. A. The role of nuclear architecture in genomic instability and ageing. Nat. Rev. Mol. Cell Biol. 8, 692–702 (2007).

103. Oberdoerffer, P. et al. SIRT1 redistribution on chromatin promotes genomic stability but alters gene expression during aging. Cell 135, 907–918 (2008).

104. Blokh, D. & Stambler, I. The application of information theory for the research of aging and aging-related diseases. Prog. Neurobiol. 157, 158–173 (2017).

105. Schumacher, B., Garinis, G. A. & Hoeijmakers, J. H. J. Age to survive: DNA damage and aging. Trends Genet. 24, 77–85 (2008).

106. Hoeijmakers, J. H. J. DNA damage, aging, and cancer. N. Engl. J. Med. 361, 1475–1485 (2009).

107. Bohr, V. A. & Anson, R. M. DNA damage, mutation and fine structure DNA repair in aging. Mutat. Res. 338, 25–34 (1995).

108. Wang, S., Prizment, A., Thyagarajan, B. & Blaes, A. Cancer Treatment-Induced Accelerated Aging in Cancer Survivors: Biology and Assessment. Cancers 13, (2021).

109. Shafqat, S., Arana Chicas, E., Shafqat, A. & Hashmi, S. K. The Achilles’ heel of cancer survivors: fundamentals of accelerated cellular senescence. J Clin Invest 132, (2022).

110. Guida, J. L. et al. Measuring Aging and Identifying Aging Phenotypes in Cancer Survivors. J. Natl. Cancer Inst. 111, 1245–1254 (2019).

111. Olsen, J. H. et al. Lifelong cancer incidence in 47,697 patients treated for childhood cancer in the Nordic countries. J Natl Cancer Inst 101, 806–813 (2009).

112. Rebholz, C. E. et al. Health care use of long-term survivors of childhood cancer: the British Childhood Cancer Survivor Study. J. Clin. Oncol. 29, 4181–4188 (2011).

113. Soto-Perez-de-Celis, E., Li, D., Yuan, Y., Lau, Y. M. & Hurria, A. Functional versus chronological age: geriatric assessments to guide decision making in older patients with cancer. Lancet Oncol. 19, e305– e316 (2018).

114. Bhakta, N. et al. Cumulative burden of cardiovascular morbidity in paediatric, adolescent, and young adult survivors of Hodgkin’s lymphoma: an analysis from the St Jude Lifetime Cohort Study. Lancet Oncol 17, 1325–1334 (2016).

115. Bullard, B. M., McDonald, S. J., Cardaci, T. D., VanderVeen, B. N. & Murphy, E. A. Nonpharmacological approaches for improving gut resilience to chemotherapy. Curr. Opin. Support. Palliat. Care 16, 151–160 (2022).

116. Liao, C.-Y., Rikke, B. A., Johnson, T. E., Diaz, V. & Nelson, J. F. Genetic variation in the murine lifespan response to dietary restriction: from life extension to life shortening. Aging Cell 9, 92–95 (2010).

117. Harper, J. M., Leathers, C. W. & Austad, S. N. Does caloric restriction extend life in wild mice? Aging Cell 5, 441–449 (2006).

118. Heilbronn, L. K. & Ravussin, E. Calorie restriction and aging: review of the literature and implications for studies in humans. Am. J. Clin. Nutr. 78, 361–369 (2003).

119. Al-Regaiey, K. A. The effects of calorie restriction on aging: a brief review. Eur. Rev. Med. Pharmacol. Sci. 20, 2468–2473 (2016).

120. Liang, Y. et al. Calorie restriction is the most reasonable anti-ageing intervention: a meta-analysis of survival curves. Sci. Rep. 8, 5779 (2018).

121. Hong, C. et al. Epigenetic Age Acceleration of Stomach Adenocarcinoma Associated With Tumor Stemness Features, Immunoactivation, and Favorable Prognosis. Front. Genet. 12, 563051 (2021).

122. Castle, J. R. et al. Estimating breast tissue-specific DNA methylation age using next-generation sequencing data. Clin. Epigenetics 12, 45 (2020).

123. Hao, J., Liu, T., Xiu, Y., Yuan, H. & Xu, D. High DNA methylation age deceleration defines an aggressive phenotype with immunoexclusion environments in endometrial carcinoma. Front. Immunol. 14, 1208223 (2023).

124. Abad, M. et al. Reprogramming in vivo produces teratomas and iPS cells with totipotency features. Nature 502, 340–345 (2013).

125. Jones, M. A. & Grotewiel, M. Drosophila as a model for age-related impairment in locomotor and other behaviors. Exp. Gerontol. 46, 320–325 (2011).

126. Ridgel, A. L. & Ritzmann, R. E. Insights into age-related locomotor declines from studies of insects. Ageing Res. Rev. 4, 23–39 (2005).

127. Jucker, M., Oettinger, R. & Bättig, K. Age-related changes in working and reference memory performance and locomotor activity in the Wistar rat. Behav. Neural Biol. 50, 24–36 (1988).

128. Siwak, C. T., Murphey, H. L., Muggenburg, B. A. & Milgram, N. W. Age-dependent decline in locomotor activity in dogs is environment specific. Physiol. Behav. 75, 65–70 (2002).

129. Moscrip, T. D., Ingram, D. K., Lane, M. A., Roth, G. S. & Weed, J. L. Locomotor activity in female rhesus monkeys: assessment of age and calorie restriction effects. The journals of gerontology series A: biological sciences and medical sciences 55, B373–B380 (2000).

130. Sallis, J. F. Age-related decline in physical activity: a synthesis of human and animal studies. Med. Sci. Sports Exerc. 32, 1598–1600 (2000).

131. Volkow, N. D. et al. Association between decline in brain dopamine activity with age and cognitive and motor impairment in healthy individuals. Am. J. Psychiatry 155, 344–349 (1998).

132. Walsh, R. N. & Cummins, R. A. The Open-Field Test: a critical review. Psychol. Bull. 83, 482–504 (1976).

133. Hollais, A. W. et al. Effects of acute and long-term typical or atypical neuroleptics on morphine-induced behavioural effects in mice. Clinical and Experimental Pharmacology and Physiology 41, 255– 263 (2014).

134. Petr, M. A. et al. A cross-sectional study of functional and metabolic changes during aging through the lifespan in male mice. Elife 10, (2021).

135. Chondronasiou, D. et al. Deciphering the roadmap of in vivo reprogramming toward pluripotency. Stem Cell Reports 17, 2501–2517 (2022).

136. Tinsley, F. C., Taicher, G. Z. & Heiman, M. L. Evaluation of a quantitative magnetic resonance method for mouse whole body composition analysis. Obes. Res. 12, 150–160 (2004).

137. Taicher, G. Z., Tinsley, F. C., Reiderman, A. & Heiman, M. L. Quantitative magnetic resonance (QMR) method for bone and whole-body-composition analysis. Anal. Bioanal. Chem. 377, 990–1002 (2003).

138. Champy, M., Argmann, C. A., Chambon, P. & Auwerx, J. Exploration of metabolic and endocrine function in the mouse. Standards of mouse model phenotyping 109–133 (2006).

139. Yang, G. et al. Central role of ceramide biosynthesis in body weight regulation, energy metabolism, and the metabolic syndrome. Am. J. Physiol. Endocrinol. Metab. 297, E211–24 (2009).

140. Chang, Y. et al. Ablation of NG2 proteoglycan leads to deficits in brown fat function and to adult onset obesity. PLoS One 7, e30637 (2012).

141. Crawley, J. N. Behavioral phenotyping of transgenic and knockout mice: experimental design and evaluation of general health, sensory functions, motor abilities, and specific behavioral tests. Brain Res. 835, 18–26 (1999).

142. Johnson, S. A., Fournier, N. M. & Kalynchuk, L. E. Effect of different doses of corticosterone on depression-like behavior and HPA axis responses to a novel stressor. Behav. Brain Res. 168, 280–288 (2006).

143. Winters, B. D., Forwood, S. E., Cowell, R. A., Saksida, L. M. & Bussey, T. J. Double dissociation between the effects of peri-postrhinal cortex and hippocampal lesions on tests of object recognition and spatial memory: heterogeneity of function within the temporal lobe. J Neurosci 24, 5901–5908 (2004).

144. Mumby, D. G., Tremblay, A., Lecluse, V. & Lehmann, H. Hippocampal damage and anterograde object-recognition in rats after long retention intervals. Hippocampus 15, 1050–1056 (2005).

145. Berlyne, D. E. Novelty and curiosity as determinants of exploratory behaviour. British journal of psychology 41, 68 (1950).

146. Ennaceur, A. & Delacour, J. A new one-trial test for neurobiological studies of memory in rats. 1: Behavioral data. Behav. Brain Res. 31, 47–59 (1988).

147. Heyser, C. J. & Chemero, A. Novel object exploration in mice: not all objects are created equal. Behav. Processes 89, 232–238 (2012).

148. Crawley, J. N. & Paylor, R. A proposed test battery and constellations of specific behavioral paradigms to investigate the behavioral phenotypes of transgenic and knockout mice. Horm. Behav. 31, 197–211 (1997).

149. Carter, R. J., Morton, J. & Dunnett, S. B. Motor coordination and balance in rodents. Curr. Protoc. Neurosci. Chapter 8, Unit 8.12 (2001).

150. Holmes, A., Wrenn, C. C., Harris, A. P., Thayer, K. E. & Crawley, J. N. Behavioral profiles of inbred strains on novel olfactory, spatial and emotional tests for reference memory in mice. Genes Brain Behav. 1, 55– 69 (2002).

151. Bach, M. E., Hawkins, R. D., Osman, M., Kandel, E. R. & Mayford, M. Impairment of spatial but not contextual memory in CaMKII mutant mice with a selective loss of hippocampal LTP in the range of the **θ** frequency. Cell 81, 905–915 (1995).

152. Barnes, C. A. Memory deficits associated with senescence: a neurophysiological and behavioral study in the rat. J. Comp. Physiol. Psychol. 93, 74–104 (1979).

153. Paylor, R., Zhao, Y., Libbey, M., Westphal, H. & Crawley, J. N. Learning impairments and motor dysfunctions in adult Lhx5-deficient mice displaying hippocampal disorganization. Physiol. Behav. 73, 781–792 (2001).

154. Freitag, S., Schachner, M. & Morellini, F. Behavioral alterations in mice deficient for the extracellular matrix glycoprotein tenascin-R. Behav. Brain Res. 145, 189–207 (2003).

155. Crawley, J. N. What’s Wrong With My Mouse?: Behavioral Phenotyping of Transgenic and Knockout Mice. (John Wiley & Sons, 2007).

156. Castro, B. & Kuang, S. Evaluation of Muscle Performance in Mice by Treadmill Exhaustion Test and Whole-limb Grip Strength Assay. Bio Protoc 7, (2017).

157. Mager, S. R. et al. Standard operating procedure for the collection of fresh frozen tissue samples. Eur. J. Cancer 43, 828–834 (2007).

158. Naber, S. P. Continuing role of a frozen-tissue bank in molecular pathology. Diagn. Mol. Pathol. 5, 253– 259 (1996).

159. Shabihkhani, M. et al. The procurement, storage, and quality assurance of frozen blood and tissue biospecimens in pathology, biorepository, and biobank settings. Clin. Biochem. 47, 258–266 (2014).

160. Peng, T. et al. A BaSiC tool for background and shading correction of optical microscopy images. Nat. Commun. 8, 14836 (2017).

161. Goldman, D. B. Vignette and exposure calibration and compensation. IEEE Trans. Pattern Anal. Mach. Intell. 32, 2276–2288 (2010).

162. Meilă, M. Comparing clusterings by the variation of information. in Learning theory and kernel machines 173–187 (Springer, 2003).

163. Sharpee, T. O. An argument for hyperbolic geometry in neural circuits. Curr. Opin. Neurobiol. 58, 101– 104 (2019).

164. Verleysen, M. & François, D. The curse of dimensionality in data mining and time series prediction. (Springer, 2005).

165. Kuo, F. Y. & Sloan, I. H. Lifting the curse of dimensionality. Notices of the AMS 52, 1320–1328 (2005).

166. Bellman, R. Dynamic programming, 1957. A very comprehensive reference with many economic examples is (2003).

167. Ganea, O., Becigneul, G. & Hofmann, T. Hyperbolic neural networks. (2018).

